# An IL-34–IGF-1 inflammatory axis fuels KRAS-mutant lung cancer progression

**DOI:** 10.64898/2026.07.14.738492

**Authors:** Jaroslav Zak, Hui Chen, Erpei Wang, Patrick Ozark, Giuliana Mognol, Matthew D. Park, Nadine Fournier, Priyanka Chaudhary, Jingjing Hu, Ryan Shepard, Anghesom Ghebremedhin, Marc Paradise, Jason Rivera, William J. Harris, Zihan Xu, Adam Ramadan, Birkley Lim, Marco Colonna, Miriam Merad, Michele De Palma, Mark Onaitis, Judith A. Varner

## Abstract

Macrophages are innate immune cells of embryonic or adult origin with tissue specific roles in homeostasis, disease surveillance, and wound repair that can be co-opted to promote tumor growth and spread^1–11^. An understanding of the specific roles of macrophage subsets in lung tumor initiation and progression could promote new therapeutic approaches for this deadly disease. Here, we show that KRAS^G12D^ mutations in lung epithelium drive proliferation of resident, embryonically-derived alveolar macrophages, which then promote tumor cell proliferation and protection from ferroptosis, leading to tumor progression. Using genetically engineered mouse models of mutant KRAS^G12D^ non-small cell lung cancer^12,13^, we found that alveolar macrophages accumulate by proliferation in response to tumor cell-secreted IL-34, recapitulating events observed in late embryonic lung development. Tumor alveolar macrophages in turn drive IGF-1-dependent tumor cell proliferation. Neutralization or deletion of IL-34 suppresses IGF-1 expression, reduces macrophage and tumor cell proliferation and inhibits tumor progression. High *IL34* and *IGF1* correlate with poor survival in KRAS^G12D/V^ lung adenocarcinomas and in other solid tumors, indicating that bi-directional proliferative signaling between resident macrophages and tumor cells can drive human lung tumor progression. These studies identify resident macrophage-tumor cell interactions as key interception points for lung cancer therapy.

---

Lung cancer remains the leading cause of death from cancer in the United States. The most common subtype of lung cancer, lung adenocarcinoma (LUAD), accounts for 40% of all lung cancers and is diagnosed both in smokers and increasingly in never-smokers. Although therapies targeting driver mutations in EGFR, KRAS, ALK, MET and other genes, as well as immune therapies, have made remarkable advances in treating LUAD patients, the overall survival rates remain low, at 28%^14^. Understanding the microenvironmental drivers of LUAD, particularly in the earliest stages of disease, could lead to new therapeutic approaches that enhance survival from this disease. A substantial body of research has shown that macrophages can promote tumor immune suppression and progression through the expression of cytokines, chemokines, and extracellular matrix factors^8–9^. Within the lung microenvironment, specialized subsets of macrophages, including alveolar, interstitial and recruited, bone marrow-derived macrophages are spatially and functionalizing distinct; each has the potential to differentially contribute to lung tumor initiation and/or progression. Recent analysis of human and murine pre-malignant and minimally invasive lung lesions identified close, functional associations between IL1β+ macrophages and IL1R+ tumor cells, although the specific origin of these driver macrophages was not identified^15^. Studies of human and genetically engineered murine KRAS-mutant identified distinct roles for resident alveolar macrophages in early lung tumor development and for bone marrow derived macrophages in lung tumor progression, although other models of KRAS-mutant tumor development identified roles for both throughout lung tumor progression^10,16–18^. Additional studies identified key roles for CD36+ alveolar macrophages in EGFR-mutant lung tumor progression^19^. Further detailed understandings of macrophage subsets and their contributions to lung cancer progression could lead to new therapeutic approaches for this deadly disease.

## Alveolar macrophage expansion in autochthonous *Kras^G12D^* mutant lung tumors

To investigate the roles and mechanisms by which resident and recruited macrophages promote lung adenocarcinoma cancer development and progression, we first analyzed the distribution, types and density of macrophages in normal human lungs and in early stage (stage 1-2) human lung adenocarcinoma (LUAD) tumors (Fig. 1; Extended Data Fig. 1a-b). We observed more than five-fold increases in CD68+ lung macrophages in LUAD tumor specimens versus normal lung tissue specimens (Fig. 1a-b; Extended Data Fig 1a-b). Increased large, round alveolar macrophages or fusiform, invasive-type macrophages were observed in individual LUAD tumors, with some tumors exhibiting primarily alveolar macrophages (Fig. 1a; Extended Data Figure 1b) and others exhibiting primarily invasive-type macrophages (Extended Data Figure 1a). We also observed abundant macrophage accumulation in alveolar spaces in tamoxifen-inducible, genetically engineered, autochthonous KRAS^G12D^ mutant lung tumor models, including *SftpcCreER; LSL-Kras^G12^*, *CC10CreER;LSL-Kras^G12D^* and *CC10CreER;LSL-Kras^G12D^;LSL-p53^fl/fl^* mice^12,13^. In *SftpcCreER;Rosa26-fGFP;LSL-Kras^G12D^* mice, which survive until 5 weeks post-tamoxifen treatment, H&E and F4/80 staining reveal that lungs showed extensive inflammation consisting of accumulation of large, foamy F4/80+ macrophages in alveolar spaces surrounding adenocarcinoma nodules between two and five weeks post-tamoxifen treatment (Fig. 1c-e; Extended Data Fig. 1c-d). Similar extensive tumor nodule development is accompanied by progressive macrophage accumulation over time in alveolar spaces in lungs of *SftpcCreER; LSL-Kras^G12^*, *CC10CreER;LSL-Kras^G12D^* and *CC10CreER;LSL-Kras^G12D^;LSL-p53^fl/fl^* mice, which survive 18 weeks and 13 weeks post-tamoxifen (Fig. 1c-e; Extended Data Fig. 1c-d). Increases in lung weight accompanied tumor and inflammation development in these models (Figure 1f). The majority of macrophages in these tumor models co-expressed the mouse macrophage marker F4/80, as well as the alveolar macrophage markers SiglecF, CD11c and the resident macrophage marker MerTK, indicating that they are resident alveolar macrophages (Fig. 1g; Extended Data Figure 1e). At high magnification, H&E images of *CC10CreER;LSL-Kras^G12D^;LSL-p53^fl/fl^* lungs reveal large round, dividing foamy macrophages in alveolar spaces with characteristic central nuclei (Fig. 1g). To confirm that alveolar macrophages were increasing by proliferation, we stained tumors with antibodies directed against the macrophage marker F4/80 and the proliferation marker Ki67. We observed that up to thirty percent of tumor macrophages co-expressed Ki67, suggesting that these cells were accumulating via proliferation (Fig. 1h-i). To evaluate the physical association between tumor cells and alveolar macrophages in these models, we first performed immunostaining for GFP to confirm that only tumor cells express GFP (Extended Data Fig. 1f) and then performed dual staining for tumor cell GFP (green) and F4/80+ macrophages (red). Tumor cells and alveolar macrophages were intermingled in close association with each other in small, early lesions, while alveolar macrophages were clustered at the periphery of tumors in adenocarcinomas, as previously described^10^ (Fig. 1j; Extended Data Fig. 1g).

**Figure 1:**
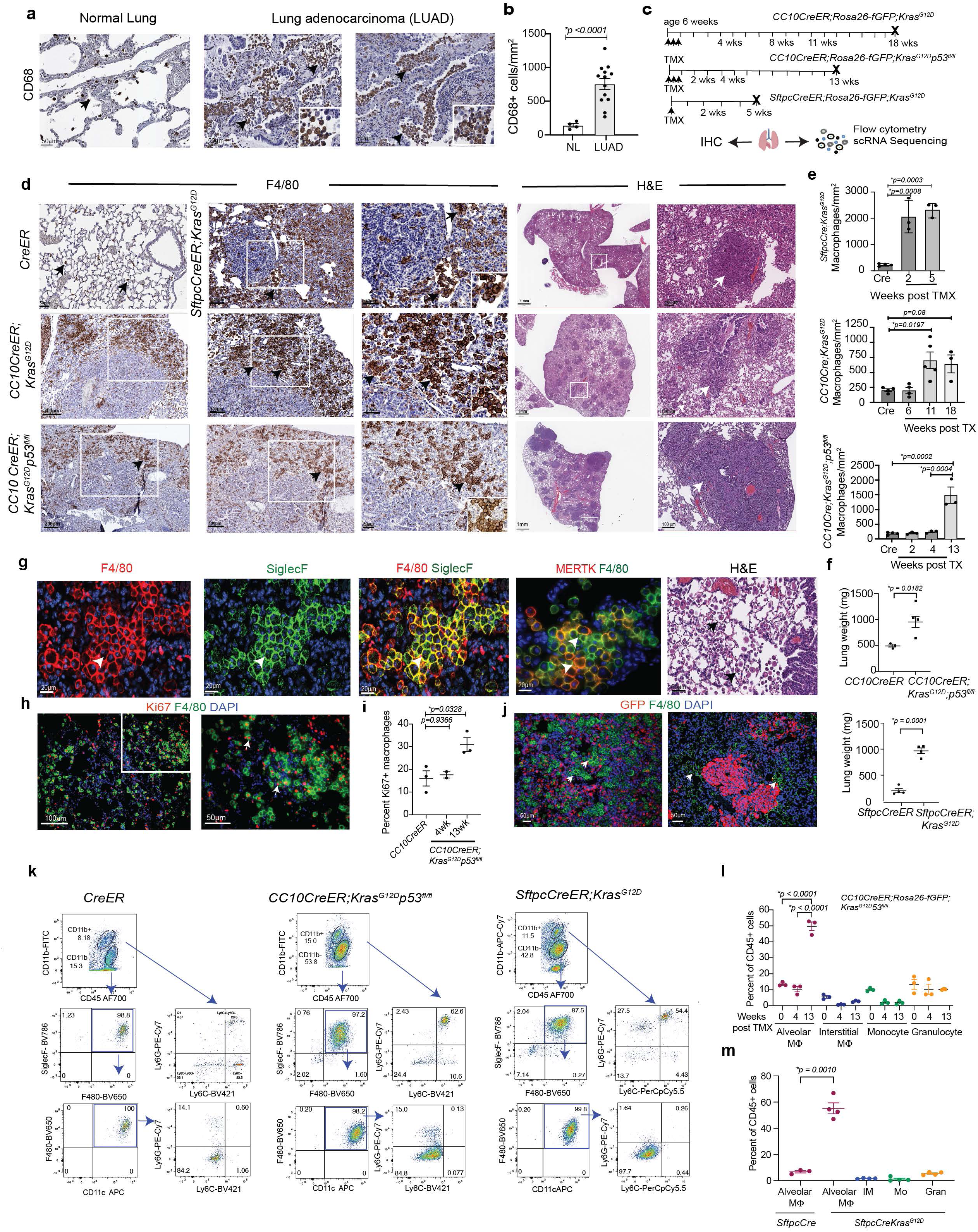
Proliferative alveolar macrophages but not bone marrow derived myeloid cells accumulate in KRAS^G12D^ mutant lung tumors. (a) CD68+ macrophages (arrows) in human normal lung and early-stage lung adenocarcinoma (LUAD). Insets: alveolar macrophages. (b) Mean CD68+ cells/mm^2^ +/− SEM in normal tissues from a (n=4-13). (c) *CC10CreER;LSL-Kras^G12D^, CC10CreER;LSL-Kras^G12D^p53^fl/fl^,* and *SftpcCreER;LSL-Kras^G12D^* mouse models of lung carcinoma: lungs were collected for analysis by histology/immunohistochemistry (IHC), flow cytometry and scRNAseq at the indicated time points until the last survival point for each model, indicated by X. (d) Images of F4/80+ macrophages (arrows) and tumors (arrows) at successive magnifications (boxes) in lungs from *CreER, SftpcCreER;LSL-Kras^G12D^*, *CC10CreER;LSL-Kras^G12D^, CC10CreER;LSL-Kras^G12D^p53^fl/fl^* animals collected at the final post-tamoxifen timepoint. Insets: alveolar macrophages. (e) Mean F4/80+ macrophages/mm^2^ +/− SEM in mouse lungs from d (n=3-5). (f) Mean +/− SEM lung weights (n=4) in mice from d. (g) F4/80+ (red), SiglecF+ (green), and F4/80+/SiglecF+ (yellow) macrophages (arrows) in lungs from *SftpcCreER;Rosa26-fGFP;LSL-Kras^G12D^* mice, as well as MerTK+(red)/F4/80+(green) (arrows) and H&E-stained macrophages in lungs from *CC10CreER;Rosa26-fGFP;LSL-Kras^G12D^* mice. (h) Images of Ki67+(red)/F4/80+(green) macrophages (arrows) in lungs from *CC10CreER;Rosa26-fGFP;LSL-Kras^G12D^p53^fl/fl^* mice. (i) Percent Ki67+ macrophages in *CC10Cre* versus *CC10Cre;LSL-Kras^G12D^p53^fl/fl^* lungs at 4 and 13 weeks post-tamoxifen (n=2-3 per timepoint, mean+/−SEM). (j) Representative patterns of GFP+ tumor cells (red) and F4/80+ macrophages (green) in lungs from developing (left) and established (right) tumors. (k-l) Flow cytometric analysis of myeloid cells in (k) *CC10CreER and CC10CreER;Kras^G12D^p53^fl/fl^* lungs 13 weeks post-tamoxifen and (l) in *SftpcCre* and *SftpcCre;LSL-Kras^G12D^* lungs 5 weeks post-tamoxifen. (m-n) Quantification of myeloid cell populations in (m) *CC10Cre* and *CC10Cre;Kras^G12D^p53^fl/fl^* lungs (n=3) and (n) *SftpcCre* and *SftpcCre;LSL-Kras^G12D^* mice (n=4). Significance determined by t-test for two groups and Anova with Tukey’s post-hoc significance testing for 3 or more groups.

In order to identify immune cells in lungs from *CC10Cre*, *SftpcCre*, *CC10Cre;Kras^G12D^;p53^fl/fl^* and *SftpcCre;Kras^G12D^* mice, we performed flow cytometry to detect myeloid cell markers of tumor-bearing and normal control lungs AMs (Extended Data Figure 2; Figure 1k-m). CD11b-Ly6C-Ly6G-F4/80+SiglecF+CD64+CD206+ alveolar macrophages were the dominant cell population in normal lungs, and this population increased by nearly five-fold in tumor-bearing lungs compared to tumor-free lungs (Fig. 1j-m). In contrast to other models, we observed decreased accumulations of Ly6C+ monocytes and Ly6C+F4/80^low^ interstitial macrophages throughout tumor development in these models of autochthonous lung tumors (Figure 1k-m). These results indicate that in proliferative, resident AMs can robustly accumulate in association with tumor cells in the lungs of autochthonous KRAS^G12D^ lung tumors; in these models, other myeloid cell populations do not significantly accumulate.

Seeking to characterize the tumor microenvironment in GEMM lung tumor models in more detail, we performed scRNA-seq of *CC10CreER* and *CC10CreER;Kras^G12C^;p53^fl/fl^* mouse lungs, near end stage at thirteen weeks after tamoxifen-treatment (Fig. 1c; 2a-d; Extended Data Fig. 3a-b). UMAPs from normal and 13 weeks post-TMX tumor-bearing lungs revealed that alveolar macrophages (AMs) were the most abundant cell type in tumor-bearing lungs (Cluster C0); this cluster was offset from and thus transcriptionally related, but distinct, from normal lung alveolar macrophages (Cluster C1) (Fig. 2a-d, Extended Data Fig. 3a-b). Fewer monocytes (Cluster C4) and T cells (Cluster C2) but more FoxP3+ regulatory T cells as well as greater T cell expression of *Tox*, a transcription factor induced in chronically stimulated, exhausted T cells, characterized tumor-bearing versus normal lungs (Fig. 2c, Extended Data Fig. 3c-e). AMs in normal and tumor-bearing lungs were defined by expression of the common AM markers *Itgax*, *Siglecf*, *Pparg*, *Cd81, Cd36, Fabp4, Mcemp1, and Il18* and by the absence of bone marrow derived monocyte/macrophage (BMDM) markers, such as *Ly6c2, Ccr2* and *Plac8* or interstitial-derived macrophage markers *C1qa*, *Selenop* and *Folr2* (Fig. 2d; Extended Data Fig. 3b). AMs of normal and tumor-bearing lungs were distinguished by unique markers, with normal AMs expressing higher levels of *Ear1, Fabp1, Flt1,* and *Kcnip4,* and tumor AMs expressing higher levels of *Gpnmb, Trem2, Igf1, Wfdc17, Fabp5*, and *Fabp4* (Fig. 2e; Extended Data Fig. 3e). AMs in tumors were characterized by signatures of oxidative phosphorylation, adipogenesis, fatty acid metabolism and MTORC1 signaling, while normal AMs expressed signatures of inflammation relative to tumor AMs; monocytes (C4) were characterized by signatures of inflammation and an absence of oxidative phosphorylation (Extended Data Fig. 3f). MKI67+ tumor AMs (C7) were uniquely characterized by signatures of proliferation (E2F Targets, G2M Checkpoint, Mitotic Spindle) (Extended Data Fig. 3f). To determine whether tumor AMs (C0) arise from normal AMs (C1), proliferating AMs (C7), interstitial macrophages (C10) or monocyte/macrophages (C4), we performed RNA velocity analysis^20–21^ of normal and tumor-bearing lungs (Fig. 2f). These analyses predicted that tumor AMs develop from MKI67+ AMs and not from monocytes or interstitial macrophages; furthermore, tumor AMs represent an immature state of normal AMs as they express reduced levels of normal AM genes and are positioned as transitional states between *Mki67+* proliferating AMs and normal AMs (Fig. 2b, 2e-f).

**Figure 2:**
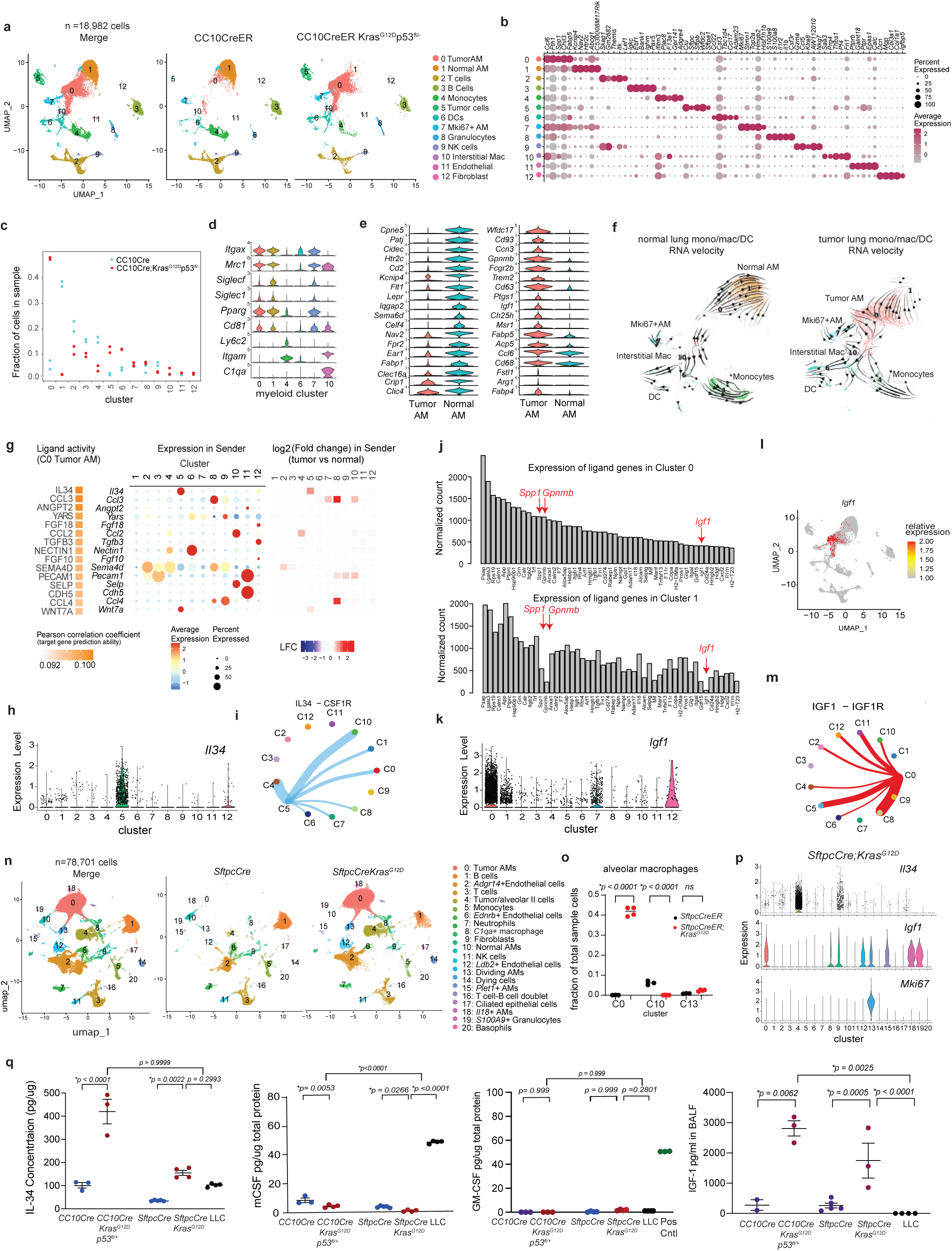
A bi-directional IL34- and IGF1-mediated proliferative signaling axis develops between resident macrophages and KRAS^G12D^ mutant lung epithelium. (a) UMAP plots from scRNA-seq of lungs from *CC10CreER and CC10CreER;Kras^G12D^p53^fl/fl^* animals 13 weeks after tamoxifen administration. (b) Dot plots of population-defining markers, (c) proportions of cell populations and (d) violin plots of monocyte/macrophage marker expression in myeloid cell clusters in *CC10CreER and CC10CreER;Kras^G12D^p53^fl/fl^* samples. (e) Violin plots of top markers expressed in normal and tumor AM. (f) RNA velocity plots of normal and tumor myeloid cell clusters from a. (g) NicheNet prediction of ligand activity and incoming cluster signaling in TAAMs (C0): (Left) Pearson correlation coefficient of top scoring ligands, (middle) expression of ligands in sender clusters, and (right) relative expression of ligands in tumor vs normal sender clusters. (h) Violin plot of *Il34* expression in clusters from a. (i) CellChat network plot of IL-34-CSF1R signaling in clusters from a. (j) NicheNet-annotated tumor cell (cluster 5) ligands expressed in normal AMs (Cluster 1) and TAAMs (Cluster 0). Differentially expressed ligands identified by red arrows. (k) Violin plot of *Igf1* expression in clusters from a. (l) Feature plot illustrating expression of *Igf1* in TAAMs. (m) CellChat network plot showing IGF1-IGF1R signaling in clusters from a. (n) Merged and groupwise UMAPs from *SftpcCreER* and *SftpcCreER;Kras^G12D^* animals. (o) Proportion of TAAMs (C0), AMs (C10) and Ki67+ AMs (C13) in *SftpcCreER* and *SftpcCreER;Kras^G12D^* animals. (p) Violin plots of *Il34, Igf1* and *Mki67* expression in clusters from n. (q) IL-34, mCSF, and GM-CSF in lung lysates and IGF-1 expression in BALF from *SftpcCreER*, *SftpcCreER;Kras^G12D^*, *CC10CreER, and CC10CreER;Kras^G12D^p53^fl/fl^* animals and in LLC conditioned medium. Significance determined by one-way ANOVA with Tukey’s posthoc correction (o,q).

## IL34-IGF1 macrophage-tumor cell signaling in *KrasG12D* mutant lung tumors

As identification of potential receptor/ligand pairs in *CC10CreER;Kras^G12D^;p53^fl/fl^* tumors that promote AM accumulation and/or proliferation could lead to new targets for lung cancer immune therapy, we used NicheNet^22^, a publicly available tool to assess ligand-receptor interactions with a large database of cytokine target genes, as well as a predictive model of ligand activity derived from single-cell transcriptomic profiles. These studies unexpectedly identified IL-34, a ligand for the myeloid cell receptor CSF1R^23–25^, as the major cytokine with ligand activity toward tumor AMs (C0) that is upregulated by KRAS^G12D^ tumor cells (C5) (Fig. 2g; Extended Data Fig. 3g). While IL-34 is required for normal development and homeostasis of microglia and Langerhans cells, resident macrophages of the CNS and epidermis, respectively, it has not been reported as an essential factor for lung macrophage development or homeostasis^23–25^. Notably, NicheNet analyses identified *Atp6v0d2*, an essential regulator of macrophage phagocytosis and cholesterol efflux, as a key downstream intermediate of the IL-34-CSF1R pathway that is upregulated in tumor AMs^26–28^, consistent with GSEA analyses of tumor AM gene expression (Extended Data Fig. 3f).

While *Il34* was strongly expressed by mutant KRAS tumor cells (C0), *Csf1* and *Csf2* were not expressed by tumor cells (Fig. 2h; Extended Data Fig. 3h). Tumor AMs and neutrophils expressed low levels of *Csf1*, *Csf2* was poorly expressed in any cells (Extended Data Fig. 3h). Accordingly, the mRNA for CSF1 receptor, *Csf1r*, was expressed in AMs, with a mean expression about half of that expressed in monocytes (C4) and interstitial macrophages (C10) (Extended Data Fig. 3i). These results were surprising, as GM-CSF, encoded by *Csf2*, is required for normal development of AMs and for AM proliferation in EGFR mutant lung cancer models^29–30^.

We next applied the CellChat v2 algorithm^31^ to infer intercellular communication networks between tumor cells and other clusters in *CC10CreER;Kras^G12D^;p53^fl/fl^* lung tumors. This analysis confirmed that the tumor cell cluster (C5) in *CC10CreER;Kras^G12D^;p53^fl/fl^* lung tumors was the only significant source of IL-34, which selectively signaled to AM clusters C0, C1, and C7 (Tumor AMs), C4 (Mo/Macs), C10 (interstitial macrophages) and other myeloid cells (Fig. 2i).

To infer active ligands expressed by AMs that could signal to tumor cells, we used the NicheNet-derived database of 643 ligands^22^. We identified *Igf1*, *Spp1*, and *Gpnmb* as ligands for receptors expressed by tumor cells that are upregulated in tumor AMs compared to normal AMs (Fig. 2j). Violin and feature plots confirmed *Igf1* upregulation in tumor AMs (C0 and C7) (Fig. 2k-l; Extended Data Fig. 3j). CellChat v2 analysis demonstrated that tumor AMs (C0) and proliferating AMs (C7) were predicted to signal via IGF-1 to IGF-1R-expressing tumor cells and granulocytes and that tumor AMs are the dominant sender of this cytokine (Fig. 3m; Extended Data Fig. 3k-m). Prior work implicates IGF-1/IGF-1R signaling in lung cancer progression through complementary mechanisms, including IGF1R-dependent PI3K activation in KRAS-mutant lung cancer cells, enhanced lung cancer stem-like phenotypes, and modulation of the lung microenvironment^32–35^. Taken together, these results suggest that bidirectional IL-34-IGF1 signaling fuels self-perpetuating lung cancer progression by promoting proliferation, survival and transcriptional modifications of AMs and tumor cells.

**Figure 3:**
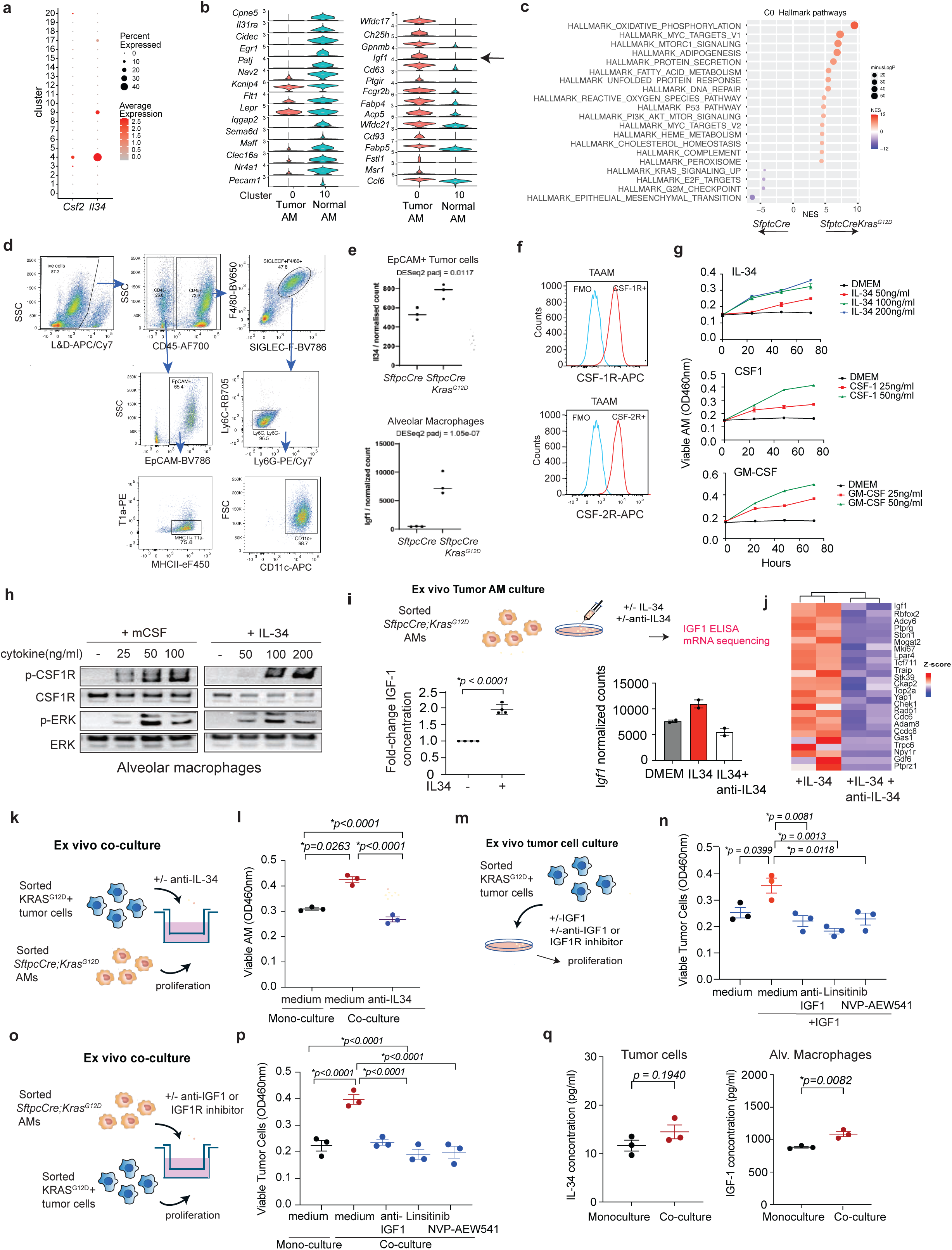
IL-34 and IGF1 promote proliferation of SpcCre;LSL-Kras^G12D^ alveolar macrophages and tumor cells. (a) Dot plots of *Csf2* and *Il34* expression in clusters from *SftpcCre;Kras^G12D^* mice. (b) Violin plots of top transcripts, including *Igf1*, that are differentially expressed in normal versus tumor AM from *SftpcCre* and *SftpcCre;Kras^G12D^* mice. (c) GSEA Hallmark pathway expression in tumor versus normal AM from *SftpcCre* and *SftpcCreER;Kras^G12D^* lungs. (d) Flow sorting of AMs and tumor cells from *SftpcCreER* and *SftpcCreER;Kras^G12D^* mice. (e) Normalized counts of *Igf1* and *II34* genes in sorted tumor AMs and tumor cells (n=3). (f) CSF1R and GM-CSF1R expression on tumor AMs, as determined by flow cytometry (n=3). (g) Proliferation of flow-sorted tumor AMs in the presence of various concentrations of IL-34, CSF1 or GM-CSF (n=4). (h) CSF1R and ERK1/2 tyrosine phosphorylation in AMs stimulated ex vivo with CSF1 or IL-34. (i) Schematic of AM culture in the presence and absence of IL-34 with or without anti-IL34; concentration of IGF-1 (n=4) in AM culture supernatant; and normalized *Igf1* counts (n=2) in cultured AM bulk sequenced RNA. (j) Heatmap of differentially expressed genes from AMs cultured in the presence of IL-34 with or without anti-IL34. (k-l) Schematic (k) and graph (l) of sorted tumor AM viability after monoculture and co-culture with sorted tumor cells in the presence and absence of anti-IL-34 (n=3). (m-n) Schematic (m) and graph (n) of sorted tumor cell proliferation after culture in the presence of IGF1 with and without anti-IGF1 or the IGF1R inhibitors Linsitinib or NVP-AEW541 (n=3). (o-p) Schematic (o) and graph (p) of sorted tumor cell proliferation after co-culture with sorted tumor AM in the presence and absence of anti-IGF-1 or the IGF1R inhibitors Linsitinib or NVP-AEW541 (n=3). (q) Expression of IGF-1 by ex vivo cultured AMs and IL-34 by ex vivo cultured tumor cells in monoculture and co-culture (n=3). Statistical comparisons were performed by Student’s t-test (I, q) or by one-way ANOVA with Tukey’s posthoc correction (l, n, p).

To confirm the role of AMs in lung tumor development, we isolated cells by bronchoalveolar lavage (BAL) from the lungs of 8-10 week old *SftpcCre* and *SftpcCre;Kras^G12D^* animals and performed bulk RNA sequencing (Extended Data Fig. 4a-i). While BAL cells from control animals were characterized by signatures of airway cells and neutrophils, BAL cells from tumor-bearing animals were dominated by signatures of AMs as well as signatures of adipogenesis and oxidative phosphorylation (Extended Data Fig. 4a-i). Tumor-bearing BAL cells were characterized by tumor AM genes such as *Fstl1*, *Ccn3*, *Igf1*, and *Fabp5* (Extended Data Fig. 4f-g).

We next performed scRNA-seq of lungs from *SftpcCre* and *SftpcCre;Kras^G12D^* animals and observed large accumulations of tumor AMs (C0) and ATII cells/tumor cells (C4) as well as decreased normal AMs (C10) in tumor-bearing animals (Fig. 2n-o; Extended Data Figure 4j-k). As in the *CC10Cre;Kras^G12D^* model, tumor cells (C4) expressed *Il34*, while proliferating AMs (C13) expressed *Mki67.* Similar to the *CC10CreER;Kras^G12D^;p53^fl/fl^* model, tumor AMs (C0) expressed *Igf1* (Fig. 2p). ELISA assays to detect IL-34, M-CSF, GM-CSF and IGF1 in lung tissues revealed that IL-34 and IGF1 proteins were both expressed in *SfptcCre;Kras^G12D^* and *CC10Cre:Kras^G12D^* tumor-bearing lungs but not in normal lungs (Fig. 2q). Neither M-CSF nor GM-CSF protein were expressed in lungs of tumor-bearing animals, mirroring RNA seq results. Accordingly, tumor cells expressed substantial *Il34* mRNA, but *Csf2* mRNA was barely detected in tumor bearing lungs (Fig. 3a).

Similar to the *CC10Cre;Kras^G12D^;p53^fl/fl^* mouse model, tumor AMs (cluster 0) downregulated normal AM genes and upregulated tumor AM genes, including *Fabp4, Fabp5*, *Gpnmb* and *Igf1* that we observed are expressed in *CC10Cre;Kras^G12D^;p53^fl/fl^* AMs (Figure 3b). Tumor AMs exhibited signatures of oxidative phosphorylation, adipogenesis and fatty acid metabolism, similar to tumor AMs in the *CC10Cre;Kras^G12D^;p53^fl/fl^* model (Fig. 3c). To confirm these findings, we sorted AMs and ATII/tumor cells from *SftpcCre* and *SftpcCre;Kras^G12D^* animals (Fig. 3d) and performed bulk RNA-seq (Extended Data Fig 4l-p). Sorted tumor AMs expressed tumor AM marker genes and signatures of fatty acid metabolism and T cell suppression (Extended Data Fig. 4l-m), while tumor cells exhibited markers and signatures of cell cycle progression, epithelial-mesenchymal transition and *Il6-Jak-Stat3* pathway activation (Extended Data Fig. 4n-p). Bulk sequencing of sorted cells confirmed that *Igf1* was upregulated in sorted tumor AMs, while *Il34* was upregulated in sorted tumor cells (Fig. 3e).

To confirm whether *CC10CreER;Kras^G12D^;p53^fl/fl^* tumor AMs were derived from self-renewing, lung resident macrophages rather than recruited bone marrow derived myeloid cells, we compared datasets from *CC10Cre* and *CC10CreER;Kras^G12D^;p53^fl/fl^* animals to previously published data from *Kras^G12D^*;*p53*-deficient lung tumors implanted in *Map17^CreER/+^R26tdTom* mice, in which cells derived from adult hematopoietic stem cells (HSCs) express tdTomato^10^. In these studies, tdTomato+ HSC-derived cells and tdTomato-cells were separated and sequenced. Integration of the *Map17CreER*-traced dataset with the *CC10CreER;Kras^G12D^;p53^fl/fl^* dataset produced alignment within major cell types and identified common MoMac (C0) and AM (C1) clusters (Extended Data Fig. 5a-f). There was a near-complete overlap of normal lung AMs from the normal *CC10CreER* and *Map17CreER* datasets (Extended Data Fig. 5g), while *CC10CreER;Kras^G12D^;p53^fl/fl^* tumor AMs clustered between normal AMs and interstitial macrophages (C5) yet retained expression of key AM markers (Extended Data Fig. 5g-i). The original *CC10CreER;Kras^G12D^;p53^fl/fl^* clusters and the integrated clusters showed a dominant overlap between tumor AMs (*CC10CreER;Kras^G12D^;p53^fl/fl^* Cluster 0) and integrated AMs (integrated Cluster 1) and a minor overlap with the interstitial macrophage-enriched integrated Cluster 5 (Extended Data Fig. 5h). Importantly, the AM cluster C1 exhibited negligible contribution from tdTomato-labeled cells, indicating that AMs were not derived from bone marrow-derived cells during the 6-month duration of the *Map17CreER* experiment (Extended Data Fig. 5g-k). These results indicate that *CC10CreER;Kras^G12D^;p53^fl/fl^* autochthonous tumors are largely tissue resident in both normal and tumor lung.

To determine whether virally transduced mouse models of mutant KRAS driven-lung cancer are also associated with IL-34-associated increases in alveolar macrophages, we integrated single cell sequencing datasets of lungs from *SftpcCreKras^G12D^* and Cre recombinase-lentivirus infected *Kras ^LSL-G12D/+^;p53^fl/fl^* mice that were treated with vehicle or Cisplatin+anti-PD-1^18^. In both the lentivirus-induced and *SftpcCreKras^G12D^* lung tumor models, we observed increased AMs (cluster 0) that were suppressed by cisplatin + anti-PD-1 treatment (Extended Data Fig. 5l-m). Tumor cells in both models expressed *Il34, but little Csf1* or *Csf2,* while AMs in both models expressed *Igf1* (Extended Data Fig 5n-p). Together, these data show that Kras^G12D^ mutations promote the expansion of the AM population in an IL34-dependent manner and that AMs then express substantial *Igf1*.

As a role for IL34 and its receptor CSF1R in tumor alveolar macrophage accumulation is unexpected, given the known role for GM-CSF in AM proliferation during early postnatal lung development, we evaluated the expression of CSF1R and CSF2R, the GM-CSF receptor, on tumor AMs by flow cytometry. We observed that each receptor was abundantly expressed on tumor AM (Fig. 3f). Each was equally expressed on normal and tumor AMs; however, their expression was at half of the level expressed by Ly6C+ myeloid cells (Ext. Data Fig. 6a). Additionally, we observed that IL-34, CSF1 and GM-CSF each stimulated ex vivo proliferation of sorted AMs (Fig. 3g). IL-34, as well as CSF1, also stimulated CSF1R signaling within 1-2 minutes after addition to sorted AMs and in BMDM, as determined by immunoblotting for the expression of phosphoCSF1R, phosphoERK1/2 and phosphoAKT (Fig. 3h; Extended Data Fig. 6b), indicating that IL-34 activates alveolar macrophage CSF1R to stimulate proliferative signal transduction.

To determine whether IL-34 controls AM expression of IGF-1, we cultured sorted murine AM in the presence and absence of IL-34 and measured murine IGF1 in the culture supernatants by ELISA (Fig. 3i). Exposure to IL-34 doubled the secretion of IGF-1 and increased *Igf1* mRNA expression by AM, while neutralizing anti-IL-34 antibodies reversed increases in IL-34 induced *Igf1* transcription (Fig. 3i). IL-34 stimulated, while anti-IL34 suppressed, expression of proliferation pathway transcripts in AMs in vitro, including *Igf1, Mki67, Chek1, Top2a, Rad51, Cdc6,* and *Yap* (Fig. 3j). To determine whether tumor cells directly stimulate AM proliferation in an IL-34 dependent manner, sorted tumor cells were co-cultured ex vivo with AMs in the presence and absence of anti-IL34 (Fig. 3k). Co-culture with tumor cells promoted, while anti-IL34 suppressed, AM proliferation in these conditions (Fig. 3l). To determine whether IGF-1 impacts *Kras^G12D^* lung tumor cell proliferation, sorted tumor cells were cultured ex vivo in the presence and absence of murine IGF-1 with and without added anti-IGF1 or IGF1R inhibitors (1μM Linsitinib or NVP-AEW541) (Fig. 3m). IGF-1 stimulated, while anti-IGF1 and IGF1R inhibitors, inhibited, tumor cell proliferation (Fig. 3n). Ex vivo co-culture of tumor cells with AMs similarly stimulated tumor cell proliferation, in a manner that was suppressed by anti-IGF-1 and IGF1R inhibitors (Fig 3o-p). We confirmed that ex vivo cultured tumor cells expressed IL-34 in both monoculture and in co-culture with sorted AMs, while ex vivo cultured AMs expressed more IGF-1 when co-cultured with tumor cells (Fig. 3q). Together, these findings show that tumor cell intrinsic IL-34 stimulates AM expansion and promotes the expression of IGF-1, which then stimulates IGF1R-dependent tumor cell proliferation.

## IL-34 inhibition suppresses *Kras^G12D^* mutant lung tumor progression

To query whether tumor derived IL-34 promotes AM accumulation and thereby mutant KRAS lung tumor progression, we investigated the effects of targeting IL-34 with neutralizing anti-IL34 antibodies in *SfptcCre;Kras^G12D^* mice and by systemic IL-34 deletion in *SfptcCre;Kras^G12D;^Il34−/−* mutant mice (Fig. 4a). IL-34 neutralization and deletion each suppressed lung tumor progression as determined by lung weight, although antibody administration more potently inhibited lung tumor weight than did IL-34 deletion (Fig. 4b). IL-34 antibody administration and *Il34* deletion inhibited AM accumulation as measured by flow cytometry (Fig. 4c-d; Extended Data Fig. 6c) with associated increases in CD4+ and CD8+ T cells (Extended Data Fig. 6d). Immunohistochemical analysis of macrophage content also showed that IL-34 neutralization potently suppressed, while IL-34 deletion partially suppressed F4/80+ as well as SiglecF+ macrophage accumulation in *SfptcCre;Kras^G12D^* tumors (Fig. 4e-f; Extended Data Fig. 6e).

**Figure 4.**
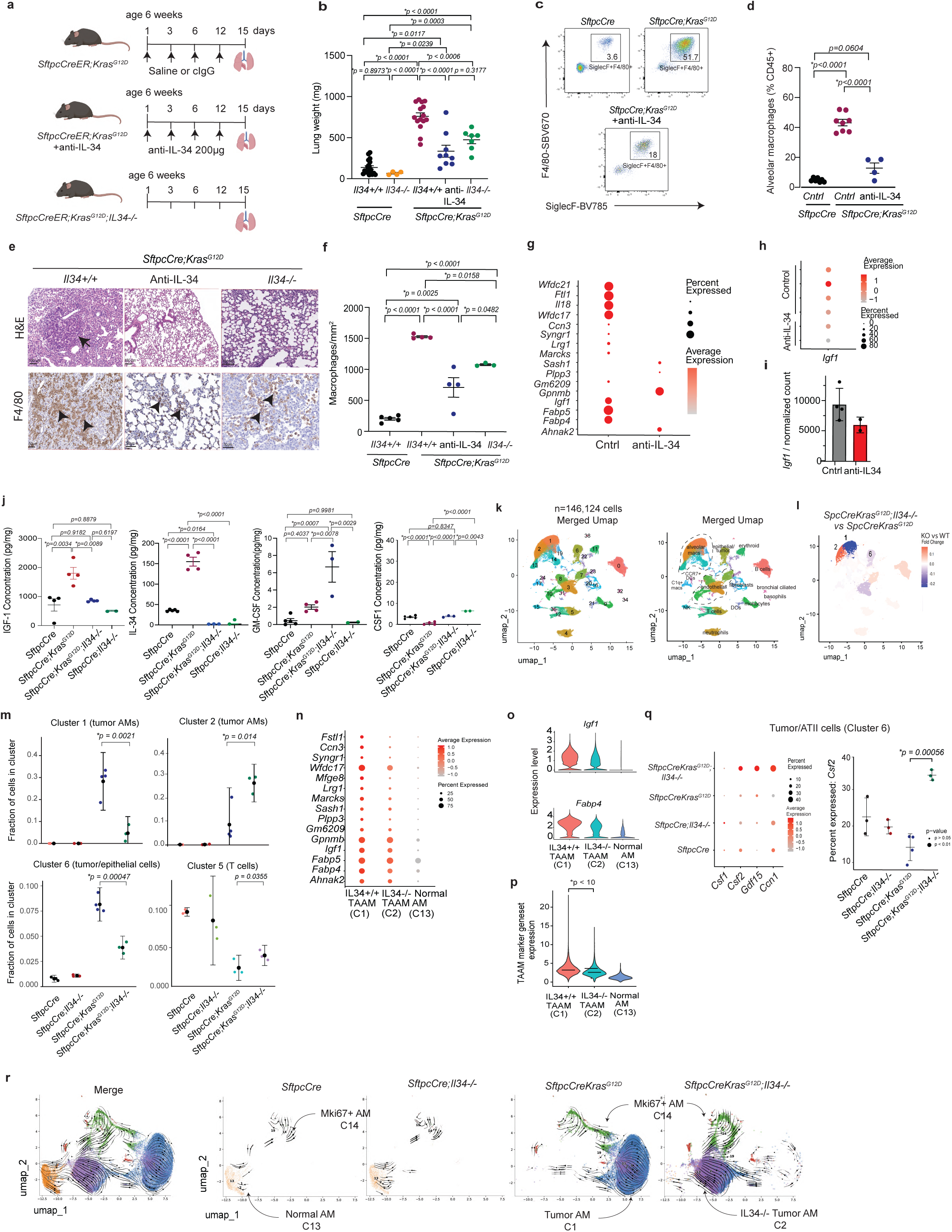
IL34 drives KRAS mutant lung tumor progression by promoting alveolar macrophage proliferation and altered transcription. (a) Schematic of anti-IL-34 treatment and *Il34* gene depletion in *SftpcCreKras^G12^*^D^ lung tumor models. (b) Lung weights from *SftpcCreER* (n=15), *SftpcCreER;Il34*−/− (n=4), *SftpcCreER;Kras^G12D^* (n=15), *SftpcCreER;KrasG12D;Il34*−/− (n=7) and *SftpcCreER;Kras^G12D^* mice treated with anti-IL34 (n=10). (c-d) Flow cytometry profiles (c) and quantification (d) of AMs in lungs from *SftpcCreER* (n=7), control treated *SftpcCreER;Kras^G12D^* (n=8) and *SftpcCreER;Kras^G12D^* mice treated with anti-IL34 (n=4). (e) H&E images and F4/80+ immunostaining to detect macrophages in lungs from *SftpcCreER;Kras^G12D^* (*Il34+/+*)*, SftpcCreER;KrasG12D;Il34*−/− and anti-IL-34 treated *SftpcCreER;Kras^G12D^* mice. (f) F4/80+ macrophages/mm^2^ in lungs from (e) (n=3-5). (g) Dot plot of differentially expressed AM genes in anti-IL-34-treated vs control-treated *SftpcCre;Kras^G12D^* animals. (h-i) Dot plot (h) and (i) graph of normalized counts of *Igf1* expression in tumor AMs from control and anti-IL-34-treated *SftpcCreKras^G12D^* animals. (j) Cytokine expression in lung lysates from *SftpcCre* (n=4), *SftpcCre;Il34*−/−, (n=2) *SftpcCreER;Kras^G12D^* (n=4) and *SftpcCreER;KrasG12D;Il34*−/− (n=3-4) mice. (k) UMAPs of scRNA-seq populations from lungs of *SftpcCre* (n=3), *SftpcCre;Il34*−/− (n=3), *SftpcCreER;Kras^G12D^* (n=4), and *SftpcCreER;KrasG12D;Il34*−/− (n=3) animals. (l) Color-coded fold-changes in the proportion of cells in clusters from *SftpcCreER;KrasG12D;Il34*−/− relative to *SftpcCreER;Kras^G12D^*lungs. (m) Fraction of cells in clusters 1, 2, 5 and 6 in each group from 4k (n=3). (n-q) Expression of (n) normal and tumor AM (TAAM) marker genes, (o) *Igf1* and *Fabp4*, and (p) normal and tumor AM marker gene signature in C1 *(Il34+/+* TAAM), C2 *(Il34−/−* TAAM) and C13 (normal AMs). (q) Expression of *Csf2* in tumor cells from treatment groups in m. (r) RNA velocity analysis of AMs in lungs from (k). Statistical comparisons were performed by one-way ANOVA with Tukey’s posthoc correction (b, d, f, j, m, p, q).

To explore the effect of IL-34 inhibition on *SfptcCre;Kras^G12D^* tumors further, we performed scRNA-seq of antibody-treated tumors. Analysis of tumors in the *SfptcCre;Kras^G12D^* anti-IL34 and control treated mice by scRNA-seq produced 16 clusters, with AMs comprising the largest cell population as well as the largest dividing cell population (Extended Data Fig. 6f-h). IL34-neutralization induced a significant shift in AMs: AMs in control-treated tumors expressed tumor-associated AM (TAAM) markers, including *Fabp4*, *Fabp5*, *Wfdc17,* and, notably, *Igf1*, while AMs in anti-IL34 treated tumors downregulated each of these markers although it had no effect on expression of *Gpnmb* (Fig. 4g-i). Furthermore, cluster 0 (C0) exhibited a 2-fold increase in relative cell abundance in anti-IL-34 treated mice, while cluster 1 (C1) was reduced almost 3-fold in the anti-IL-34 group (Extended Data Fig. 6i-j). Examination of transcripts distinguishing C0 from C1 identified TAAM markers, including *Fabp4*, *Fabp5*, *Igf1* and *Wfdc17* as significantly more highly expressed in C1 whereas C0 expressed higher levels of normal AM markers such as *Lepr* (Fig. 4g; Extended Data Fig. 6k-l). Furthermore, pseudobulk differential expression analysis of anti-IL34 vs control treated samples revealed that TAAM markers such as *Fabp4*, *Marco* and *Wfdc21* were downregulated in the anti-IL-34 group even within C0 and C1 (Extended Data Fig. 6m-n). Therefore, anti-IL34 treatment downregulated TAAM marker expression and restored normal AM marker expression in tumor-infiltrating AMs. In tumor cells (C6), there was also a significant downregulation of transcripts associated with angiogenesis and inflammation such as *Il1b* and with cancer invasiveness such as *Lgals1* and stress responsiveness, such as *Gpx1,* in the anti-IL34-treated group (Extended Data Fig. 6o). GSEA analysis of AM clusters C0 and C1 further showed that the oxidative phosphorylation, fatty acid metabolism and EMT/TGFbeta signaling pathways were downregulated and TNF-mediated pro-inflammatory pathways were upregulated in both AM clusters from anti-IL-34 treated animals (Extended Data Fig. 6p-s).

We also performed anti-IL34 and control treatment in *CC10CreER;Kras^G12D^;p53^fl/fl^* tumor bearing mice. IL-34 neutralization suppressed AM accumulation in *CC10Cre;Kras^G12D^p53^fl/^*^fl^ lung tumors as measured by MerTK+F4/80+ macrophage content (Extended Data Fig. 7a-b). scRNA-seq analysis of these tumors identified 14 clusters including AMs (Cluster 2), although the relative percentage of AMs of total cells was lower than in the *SfptcCre;Kras^G12D^* model (Extended Data Fig. 7c-e). Similar to the *SfptcCre;Kras^G12D^* anti-IL34 treatment experiment, anti-IL34-treated *CC10CreER;Kras^G12D^;p53^fl/fl^* mice exhibited lower *Igf1* expression in tumor-infiltrating AMs (Cluster 2) by differential gene expression (Extended Data Fig. 7f-g) and CellChat analyses (Extended Data Fig. 7h), indicating a loss of *Igf1* production by AMs. Fibroblasts (C6) were the only significant *Igf1* producing cluster remaining in the anti-IL34 group (Extended Data Fig. 7h). Additionally, the reduced strength of IL-34 signalling was also reflected in the reduced tumor-to-AM (C2->C5) and tumor-to-interstitial macrophage (C5->C9) signalling probability as estimated by CellChat (Extended Data Fig. 7h). These results establish that antibody-mediated IL-34 blockade in the *SfptcCre;Kras^G12D^* and *CC10CreER;Kras^G12D^;p53^fl/fl^* models results in the reduction of *Igf1* production by AMs and loss of IGF1-IGF1R signalling by AMs to other cell types.

We determined whether *Il34* deletion affected the expression of cytokines in lungs from *SfptcCre;Kras^G12D^* and *SfptcCre;Kras^G12D^*;*Il34−/−* mice by ELISA. *SfptcCre;Kras^G12D^* tumor-bearing lungs exhibited substantial upregulation of IL-34 and IGF-1 compared with non-tumor bearing control lungs; however, little CSF1 or GM-CSF was expressed in either of these samples (Fig. 4j). Substantially reduced IGF-1 and no IL-34 or M-CSF were observed in *Il34−/−* tumors. However, *SfptcCre;Kras^G12D^*;*Il34−/−* tumors exhibited a three-fold increase in GM-CSF expression compared to *SfptcCre;Kras^G12D^* tumors (Fig. 4j). These results suggest that *Il34−/−* KRAS*^G12D^* tumors partially compensate for loss of IL-34 by upregulating GM-CSF, a cytokine that promotes the early postnatal development of lung alveolar macrophages^29–30^, thus accounting for the intermediate effects of IL34-deletion compared with antibody-mediated neutralization.

We next analyzed lungs from *SfptcCre;Kras^G12D^*;*Il34−/−*, *SfptcCre;Kras^G12D^*;*Il34*^+/+^, *SfptcCre*;*Il34−/−* and *SfptcCre*;*Il34*^+/+^ mice by scRNA-seq. Dimensionality reduction and clustering of the combined dataset yielded 44 clusters of which 15 contained predominantly cell doublets and were removed from subsequent analysis (Extended Data Fig. 8a-c, Fig. 4k). The tumor samples were characterized by large tumor-enriched AM clusters (C1, C2), including a dividing AM cluster (C14), whereas AMs in normal lung samples clustered separately (C13). As in the anti-IL-34 studies, we observed a significant shift within AM subclusters: *Il34*^−/−^ tumors showed a significant reduction in C1 and significant increase in C2 (Fig. 4l, Extended Data Fig. 8b). Tumor cells (C6) were significantly reduced, and T cells (C5) were significantly enriched in *Il34*^−/−^ tumors (Fig. 4m). TAAM markers previously identified in the *CC10cre* model, including *Igf1* and *Fabp4,* were reduced in *Il34*^−/−^ vs *Il34*^+/+^ *SfptcCre* tumors (Fig. 4n-p). We analyzed the overlap between TAAM markers in the *CC10cre* and *SfptcCre* models to determine how many TAAM markers were IL-34-dependent. Of the 1,515 TAAM markers in the *SfptcCre*;Kras^G12D^ vs *SfptcCre* dataset (significantly upregulated genes in C0 vs C10, Fig. 4n), the majority (n=1,009) were also significantly upregulated in CC10Cre TAAMs (Extended Data Fig. 8d). Of these 1,009 shared TAAM markers, 687 were significantly downregulated in *Il34*^−/−^ enriched AMs (C2) compared to *Il34*^+/+^ tumor enriched AMs (C1) (Extended Data Fig. 8d). The expression of TAAM markers, including *Igf1* and *Fabp4,* in *Il34*^−/−^ tumors was intermediate between *Il34*^+/+^ tumor and normal lung AMs (Fig. 4n-p; Extended Data Fig. 8e). This analysis demonstrates that IL-34 controls the expression of the majority of TAAM markers shared between the *CC10Cre* and *SfptcCre* models and that deletion of IL-34 downregulates these markers.

We analyzed inter-cluster signaling in *Il34*^+/+^ and *Il34*^−/−^ tumors using CellChat. The *Il34*^+/+^ tumor-enriched AM Cluster 1 was identified as the dominant sender of IGF signaling, consistent with its strong expression of *Igf1* (Extended Data 8f). The second strongest senders of IGF signaling were dividing AMs (Cluster 14). However, despite the lack of IL-34 and reduction in IGF1 signalling in *Il34*^−/−^ tumors, AMs were still detectable and contained a proliferative subpopulation. Tumor cells in *Il34*^−/−^ mice significantly upregulated the transcript *Csf2* encoding GM-CSF and its receptor transcript, *Csf2ra*, as well as the stress response gene *Gdf15* and the matricellular gene *Ccn1* in tumor AMs (Fig. 4q). Cell Chat analysis showed that GM-CSF signalling by tumor cells was only significant in the *Il34*^−/−^ but not *Il34*^+/+^ tumors, supporting the conclusion that *Csf2* expression in *Il34−/−* tumors compensates for *Il34* loss (Extended Data Fig. 8g-i). As GM-CSF is required for the development of normal lung AMs, its upregulation in *Il34*^−/−^ tumors suggest these tumors compensate for the loss of IL-34 by utilizing an alternative AM growth factor. In line with the reduced tumor-associated signature of AMs in *Il34*^−/−^ mice, we also observed an upregulation of T cell activation and interferon signalling molecules such as *Stat1* and *Irf1* in *Il34*^−/−^ tumors (Extended Data Fig. 8j).

To better understand the relationship between tumor AMs, normal AMs and the effect of IL-34 on tumor AMs, cells in AM clusters from the combined scRNA-seq dataset were extracted and subjected to re-normalization and dimensionality reduction, then RNA velocity analysis. The resulting velocity vectors imply that the dividing Mki67+ AM cluster (C14) gives rise to AMs in C1 (*Il34*+/+ TAAM), C2 (*Il34−/−*TAAM) or C13 (Normal AM), with C13 being the most differentiated cluster (Fig. 4r). RNA velocity vectors were not dramatically different between *Il34*^−/−^ and *Il34*^+/+^ normal lung AMs (Fig. 4r), but *Il34*^−/−^ tumor AMs had a distinct pattern from *Il34*^+/+^ tumor AMs: *Il34*^−/−^ tumor AMs clustered closer to normal AMs and RNA velocity vectors were consistent with a direct generation of these more differentiated cells from the dividing cluster C14 (Fig. 4r). In accordance with these results, random walk analysis by CellRank with cells in clusters C1, C2 or C14 uniformly predicted that most AMs in any of these clusters were likely to reach final states in the C14 cluster (Extended Data Fig. 8k). Taken together, these results are consistent with a model where proliferation of immature AMs in KRAS mutant lung cancer is driven by IL-34 signalling. IL-34 deficiency leads to compensation by tumor-produced GM-CSF, but the resulting AMs more closely resemble normal AMs. In support of this model, bulk RNA-seq of sorted *SfptcCre;Kras^G12D^*;*Il34−/− and SfptcCre;Kras^G12D^* AMs confirmed that IL-34 deletion promotes normalization of AMs, shifting AMs, as well as tumor cells, away from oxidative phosphorylation and proliferation pathways and toward pro-inflammatory signaling (Extended Data Fig. 8l-o).

We next asked whether tumor AMs influence the functional states of tumor cells. Cells belonging to alveolar epithelial/tumor cell clusters in the *Il34*^−/−^ *SftpcCre, Il34*^++^ *SftpcCre, Il34*^−/−^*SftpcCreKras*^G12D^ vs *SftpcCreKras*^G12D^ and anti-IL34 treated *SftpcCreKras*^G12D^ vs isotype treated datasets were merged. Non-epithelial cells were filtered out, and the new dataset was subjected to re-normalization and re-clustering (Fig. 5a). Nine clusters, including large clusters of AT2-like cells (C0, C1), a dividing cell cluster (C8), and a cluster (C3) expressing stem-like markers were identified (Fig. 5a, Extended Data Fig. 9a). C3 further expressed markers of the high-plasticity cell state (HPCS), such as *Hmga2*, that were recently described in a mutant KRAS-driven tumor model^36,37^ as well as markers of the damage-associated transient progenitors (DATP) such as *Cldn4*^38^ (Fig. 5b). A transition cluster expressing AT2 markers and weakly expressing HPCS markers was noted (C2) as well as an AT1 cluster (C5) (Fig. 5a-b, Extended Data Fig. 9a). Interestingly, C1, C3 and C7 upregulated markers of cell stress, with C7 expressing a pre-ferroptosis signature including *Ddit3*, *Trib3* and *Chac1* (Fig. 5c-d, Extended Data Fig. 9b). Examination of the effects of IL-34 modulation on the relative distribution of these clusters revealed a concordant increase in the pre-ferroptotic cluster and concordant decrease in the transition and unstressed AT2-like clusters in both *Il34*^−/−^ and anti-IL34 experiments relative to respective controls (Fig. 5d, Extended Data Fig. 9c-d). Analysis of predicted cell trajectories using scvelo identified C0 as the most terminal cluster, consistent with its unstressed AT2-like signature (Fig. 5e, Extended Data Fig. 9e). Partition-based graph abstraction (PAGA) analysis predicted that cells enter this cluster predominantly from clusters C2 and C8 with a smaller contribution from C7, suggesting a resolution of the pre-ferroptotic state in a subset of cells (Fig. 5e, Extended Data Fig. 9e). Interestingly, the C7->C0 transition was predicted to be 4x less probable in *Il34*^−/−^ tumors despite C7 having a 3x higher relative abundance in *Il34*^−/−^ tumors, suggesting *Il34*^−/−^ AT2-like tumor cell struggle to exit from pre-ferroptosis (Fig. 5e, Extended Data Fig. 9f-g). Self-transition of the intermediate cluster C2 into C2 was sharply reduced in *Il34*^−/−^ (Extended Data Fig. 9g). Transitions from C8 to C3 and C2 were also predicted to be less probable, instead redirected towards C1, suggesting the actively dividing cell pool in *Il34*^−/−^ tumors preferentially produces stressed AT2-like tumor cells rather than replenishing the stem-like pool (Fig. 5e). *Il34* expression was relatively uniform across most clusters, with highest levels in the pre-ferroptotic cluster C7 (Extended Data Fig. 9h).

**Figure 5.**
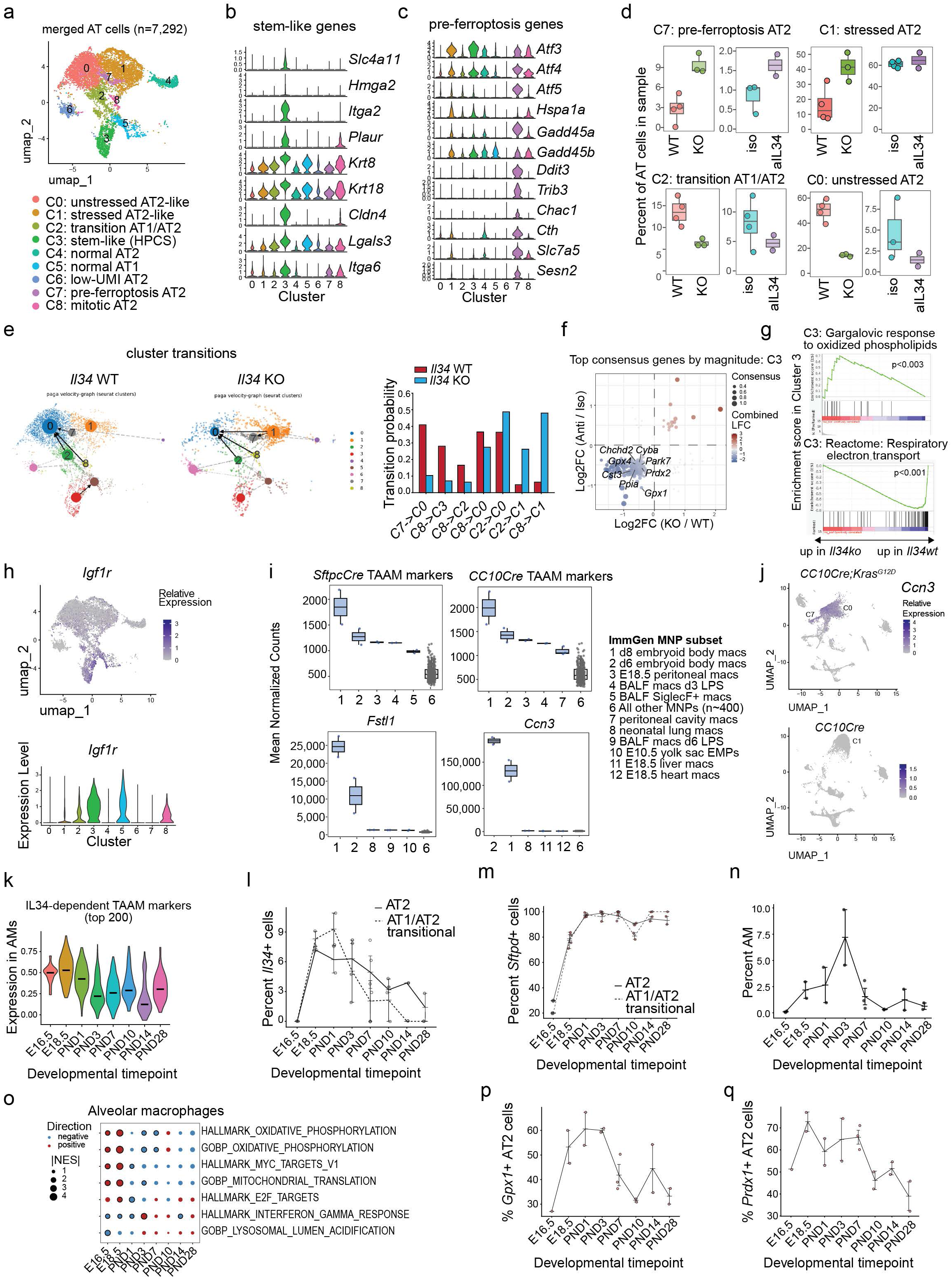
De-differentiation and oxidative stress management in KrasG12D tumor cells and tumor-associated alveolar macrophages. (a-h) scRNA-seq analysis of merged AT2 and AT1-like cells from *Il34*^−/−^ and anti-IL34 *SftpcCreKras*^G12D^ experiments: from Figure 4. (a) UMAP dimensionality reduction and clustering; (b) violin plot of stem-like (HPCS and DATP) markers; (c) violin plot of pre-ferroptosis markers; (d) box plots of relative cluster shared by experimental group; (e) partition-based graph abstraction analysis vectors in UMAP space and changes in cluster transition probabilities in *SftpcCreKras*^G12D^ *Il34*^−/−^ and *Il34*^+/+^ groups; (f) consensus differentially expressed genes between *SftpcCreKras*^G12D^ *Il34*^−/−^ and *Il34*^+/+^ cells (horizontal axis) and anti-IL34 vs control cells (vertical axis) in Cluster 3, axes show log_2_fold change; (g) gene set enrichment analysis comparison of *SftpcCreKras*^G12D^ *Il34*^−/−^ and *Il34*^+/+^ cells in Cluster 3; (h) feature plot of Igf1r expression; (i) Mean expression of TAAM marker genes, *Fstl1* and *Ccn3,* in mononuclear phagocyte subsets from ImmGen. The top 5 highest-expressing subsets are shown for each analysis followed by all other MNP subsets grouped. (j) Feature plot of *Ccn3* expression in the *CC10creKras*^G12D^ tumor and control groups. (k) Violin plot of relative expression of top IL34-dependent TAAM genes at developmental timepoints. (l-m) Percentage of *Il34*- (l) and *Sftpd*- (m) expressing AT2 and AT1/AT2 transitional cells grouped by developmental timepoint. (n) Percentage of AMs of total lung cells in each sample grouped by developmental timepoint. (o) Bubble plot of normalized enrichment scores of selected gene sets with significant changes in AM expression between developmental timepoints. Statistically significant changes are marked with a rimmed circle. (p-q) Percentage of *Gpx1*- (p) and *Prdx1*- (q) expressing AT2 cells grouped by developmental timepoint. Statistical analysis was performed using the Seurat and DESeq2 packages (f) and GSEA applied to pseudobulk data (g, o). TAAM, tumor-associated alveolar macrophages; AT2, alveolar epithelial type 2 cells; UMAP, uniform manifold approximation and projection; ImmGen, Immunological Genome Project. HPCS, high plasticity cell state. DATP, damage-associated transient progenitors.

Differential gene expression analysis identified the highest number of significant DEGs in cluster C3 including the concordant downregulation of multiple antioxidant genes in C3 in *Il34*^−/−^ and anti-IL34 treated tumors (Fig. 5f, Extended Data Fig. 9i). Genes associated with the response to oxidized phospholipids were upregulated and oxidative phosphorylation genes downregulated in *Il34*^−/−^ cells (Fig. 5g). C3 cells exhibited a striking high expression of *Igf1r*, suggesting a possible role for IGF1 signaling in maintaining gene expression in these stem-like cells (Fig. 5h). Given the upregulation of response to oxidized lipids and increase in the pre-ferroptotic cluster, we experimentally assessed oxidized lipids in *Il34*^−/−^ and *Il34*^+/+^ tumor cells, observing a significant increase in lipid oxidation in the *Il34*^−/−^group (Extended Data Fig. 9j). These results demonstrate that IL-34 is required for the maintenance of antioxidant gene expression and prevention of lipid oxidation in *Kras*^G12D^ lung tumor cells and the sustained generation of unstressed AT2-like tumor cells from dividing and stem-like pools. Given that IGF1 is a known factor for the maintenance and proliferation of stem cells in multiple tissues and tumor types including lung cancer^39–40^, it is likely that IL-34 mediated production of IGF1 is required to support HPCS self-renewal, HPCS-driven tumor growth and prevention of ferroptosis.

To better understand the transcriptomic changes in TAAMs and the role of IL-34, we examined the expression pattern of TAAM markers in the large collection of mononuclear phagocytes (MNP) by ImmGen (GSE1221080)^41^, Interestingly, the ImmGen MNP subset most highly enriched for TAAM markers were embryoid body macrophages differentiated from stem cells, followed by E18.5 embryonic peritoneal macrophages (Fig. 5i). This effect was exemplified by genes such as *Fstl1* and *Ccn3* which exhibit high expression in embryoid body, embryonic/neonatal and tumor macrophages but low-to-zero expression in normal adult AMs (Fig. 5i-j).

To corroborate this link between embryonic gene expression and TAAMs, we analyzed the publicly available scRNA-seq LungMAP dataset of the developing mouse lung (LMEX0000004397)^42^ comprising 8 developmental timepoints between E16.5 and postnatal day 28 (PND28) (Extended Data Fig. 9k). In accordance with the ImmGen MNP analysis, top TAAM markers were most highly expressed in embryonic AMs, and this effect was even more pronounced for IL-34-dependent TAAMs (Fig. 5k, Extended Data Fig. 9l). The expression of *Il34* transcript was detected in AT2 and AT1/AT2 transitional cells, peaking at E18.5-PND1 and decreasing towards adulthood (Fig. 5l, Extended Data Fig. 9m). In contrast, the expression of canonical AT2 markers such as *Sftpd* increased in the embryonic stage and stabilized from PND3 (Fig. 5m, Extended Data Fig. 9n). Interestingly, we observed a major expansion of AMs peaking at PND3 followed by a reduced % of total lung cells (Fig. 5n). The expansion period of AMs coincides with the developmental phase E18.5-PND3 during which *Il34* and *Csf2* but not *Csf1* are highly expressed in AT2 and AT1/AT2 cells (Extended Data Fig. 9o-p). Most AT2 cells expressed *Il34* or *Csf2* but not both (Extended Data Fig. 9q). Genes encoding the IL-34 and CSF2 receptors were also expressed during this phase (Extended Data Fig. 9r-s). At a pathway-level, embryonic AMs expressed significantly higher levels of genes related to oxidative phosphorylation, mitochondrial translation and MYC targets whereas genes related to interferon response or lysosomal acidification were enriched in postnatal AMs, consistent with a known proliferative, metabolically active phenotype of embryonic AMs (Fig. 5o).

Hierarchical clustering of genes expressed in AT2 cells along development identified a cluster containing *Il34* characterized by high expression at E18.5-PND1. This cluster notably contained antioxidant genes *Gpx1* and *Prdx1* (Fig. 5p-q, Extended Data Fig. 9t-v), and genes temporally co-expressed with *Il34* were significantly enriched in the Gene Ontology Response to Oxidative Stress gene set (Extended Data Fig. 9w). The HPCS marker *Hmga2* was expressed in embryonic and early postnatal but not adult AT2 cells, highlighting that the stem-like cancer cells upregulate some embryonic genes (Extended Data Fig. 9x). Taken together, these results demonstrate that embryonic gene expression programs are reactivated in the *Kras*^G12D^ tumor cells and TAAMs, partially driven by the activation of oxidative stress response and OxPhos pathways.

## IL-34 expression and IL-34-induced tumor-associated AM markers are associated with shorter survival in human KRAS-G12D/V mutant lung tumors

To determine whether the pro-tumor effects of AMs and IL-34 signaling in mouse lung cancer were relevant to human lung cancer, we re-analyzed a publicly available scRNA-seq dataset of human non-small cell lung cancer-infiltrating immune cells^17^. Comparing AM content by tumor stage confirmed the previously published enrichment of AMs in earlier-stage tumors; T1 stage tumors exhibited the highest AM content (Extended Data Fig. 10a). KRAS-mutant tumors had a higher AM content than other mutation groups (Extended Data Fig. 10b) and a significantly higher content of AMs compared to KRAS-wild type tumors (Fig. 6a). Differential expression analysis of AMs in KRAS-mutant vs KRAS-wild type tumors revealed a number of murine TAAM markers including *FABP4*, *FTL* and *IGF1* were significantly upregulated in KRAS-mutant compared to KRAS-wild type tumors (Fig. 6b-c, Extended Data Fig. 10c). Human orthologues of murine TAAM markers were significantly more highly expressed in tumor AMs than normal lung AMs (Extended Data Fig. 10d).

**Figure 6.**
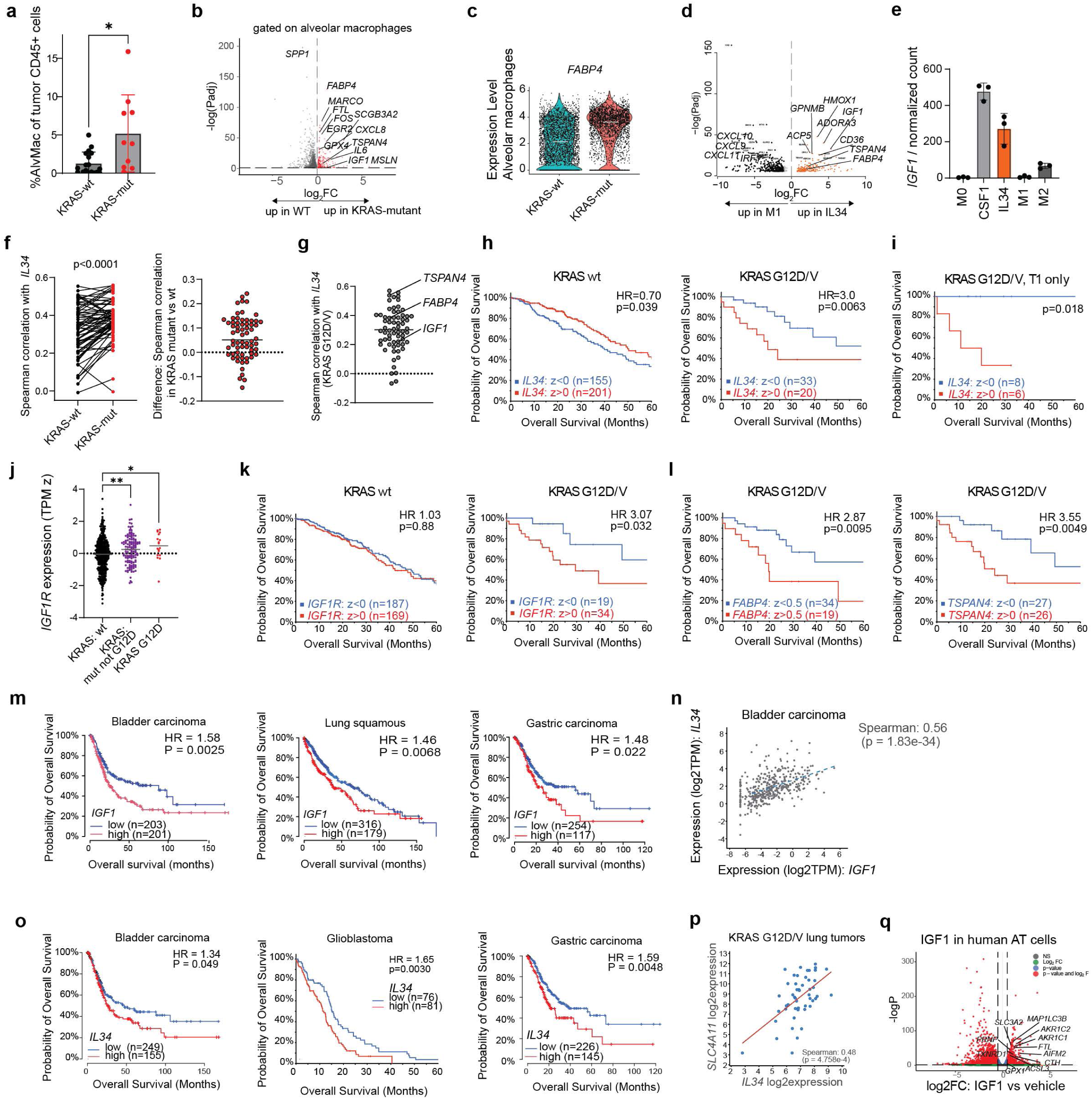
IL34-IGF1 signalling axis correlates with survival in human cancer progression. (a) Percent of tumor AMs, (b) volcano plot of differentially expressed genes in tumor AMs and (c) expression of *FABP4* in tumor AMs in KRAS-mutant vs KRAS-wild type tumors using reanalysis of publicly available data by Leader et al.^17^ (d-e) Volcano plot of differentially expressed genes (d) and normalized expression of IGF1 (e) in human monocytes treated with IL-34 vs M1 polarizing cytokines (IFNγ and GM-CSF) based on reanalysis of public dataset GSE151194. (f) Spearman correlation of 66 consensus IL34-dependent TAAM marker genes with *IL34* in KRAS-mutant and KRAS-wild type tumors in TCGA LUAD dataset (left); difference between Spearman correlation of 66 TAAM marker genes and *IL34* in KRAS-mutant vs wild type tumors (right). (g) Dot plot of absolute Spearman correlation of 66 IL34-dependent TAAM marker genes with *IL34* expression in KRAS-G12D mutant TCGA LUAD tumors with notable genes highlighted. (h) Overall survival analysis of high vs low *IL34* expression in KRAS-wt and KRAS-G12D or G12V TCGA LUAD tumors. (i) OS analysis of IL34 expression in KRAS G12D/V stage T1 tumors only (TCGA LUAD). (j) Relative expression of *IGF1R* (z-score of transcript per million) in KRAS wild type, KRAS mutant but not G12D, and KRAS G12D LUAD tumors. (k) OS analysis of high vs low *IGF1R* expression in KRAS wild type and KRAS G12D/V LUAD tumors. (l) OS analysis of *FABP4* high vs low and *TSPAN4* high vs low (right) in KRAS G12D/V LUAD tumors. (m) OS analysis of IGF1 high vs low tumors in bladder, lung squamous and gastric carcinoma TCGA datasets. (n) Correlation of *IGF1* and *IL34* expression in TCGA bladder carcinoma dataset. (o) OS analysis of *IL34* expression in TCGA solid tumor type datasets. (p) Correlation between the mRNA levels of *IL34* and HPCS marker *SLC4A11* in the TCGA lung adenocarcinoma tumors with KRAS G12D or V mutations. (q) Volcano plot of differentially expressed genes between IGF1 and control treated human AT2 cells (GSE262398); genes officially annotated as ferroptosis-modulating or antioxidant are highlighted in text. HPCS, high plasticity cell state. Statistical comparisons were performed by the log-rank test (h, i, k, l, m, o), Student’s t-test (a, j), Student’s paired t-test (f), Wilcoxon rank sum test (b) and DESeq2 Wald test (d). Bars show mean and standard deviation. OS, overall survival, TCGA, The Cancer Genome Atlas, LUAD, lung adenocarcinoma.

Re-analysis of a publicly available RNA-seq dataset of IL-34-treated human macrophages^47^ confirmed that many of these TAAM genes, including *FABP4*, *TSPAN4* and *IGF1,* are directly induced by IL-34 or CSF1 in human macrophages (Fig. 6d-e, Extended Data Fig. 10e). To assess the role of KRAS mutation status in the correlation of IL-34 with TAAM markers, the Spearman correlation coefficient between *IL34* and individual TAAM marker transcripts was calculated in KRAS-mutant and KRAS-wild type tumors. KRAS-mutant tumors exhibited significantly stronger correlation coefficients between IL-34 and TAAM markers than KRAS-wild type tumors, and virtually all these markers showed a positive correlation with *IL34* in KRAS-mutant tumors (Fig. 6f-g, Extended Data Fig. 10f).

Consensus analysis of 6 differential expression comparisons (AMs in tumor vs normal lung in *SftpcCre* and *CC10cre* models and in the Leader et al. human dataset, AMs in KRAS-mutant vs KRAS-wild type human lung cancer, IL-34-treated vs control human macrophages and AMs in *Il34^−/−^* vs wild type *SftpcCre* tumors) revealed *IGF1* and *FTL* as consensus orthologous TAAM markers dependent on IL-34 and enriched in KRAS-mutant tumors, underscoring the importance of *IGF1* as an IL-34 target (Extended Data Fig. 10g). GSEA of these 6 differential expression comparisons also showed pathway similarities: fatty acid metabolism was upregulated in tumor vs normal AMs in the *CC10cre* and *SftpcCre* mouse models as well as in human lung cancer (Extended Data Fig. 10h). Conversely, normal AMs upregulated TGF-beta signaling, which was reduced in *Il34*^−/−^ tumor AMs compared to *Il34*^+/+^ tumor AMs (Extended Data Fig. 10i). Oxidative phosphorylation was upregulated in mouse tumor AMs, *Il34*^+/+^ AMs and in human macs treated with IL-34 (Extended Data Fig.10j).

We next asked whether the expression of *IL34* had any prognostic significance in human NSCLC. Utilizing the publicly available TCGA lung adenocarcinoma dataset, tumors expressing higher levels of *IL34* had a more favorable prognosis in patients with KRAS-wild type tumors (Fig. 6h). However, this was not the case in KRAS-mutant disease, and patients with high *IL34* and *KRAS*^G12D^ or *KRAS*^G12V^ mutant tumors had a significantly shorter overall survival (OS) compared to *IL34*-low tumors in the same mutant group (HR=3.0, Fig. 6h). This effect was the most pronounced in stage T1 disease where the only observed deaths during follow up were *IL34*-high patients (Fig. 6i). Given the ability of IL-34 to induce IGF1 expression, we assessed the expression of *IGF1R* in the same dataset and found that KRAS-mutant tumors expressed significantly higher levels of *IGF1R* than KRAS-wild type tumors, with *KRAS*^G12D^ tumors expressing the highest *IGF1R* levels (Fig. 6j). IGF1R expression had no significant association with OS in KRAS-wild type cancers but was associated with significantly shorter OS in *KRAS*^G12D^ or *KRAS*^G12V^ cancers (HR=3.07, Fig. 6k) but not tumors with *KRAS* mutations other than G12D or G12V (Extended Data Fig. 10k). Other IL-34-dependent TAAM markers such as *FABP4* or *TSPAN4* similarly showed significant associations between higher expression and shorter OS (Fig. 6l). Interestingly, expression of the IGF1R ligand *IGF1* was also associated with shorter survival in other cancers, including lung squamous cell carcinoma, bladder and gastric carcinomas (Fig. 6m), and *IGF1* expression closely correlated with *IL34* expression (Fig. 6n). High *IL34* expression also showed significant associations with shorter OS in other TCGA datasets including bladder, renal, gastric, ovarian, hepatocellular, adrenocortical carcinomas and glioblastoma (Fig. 6o, Extended Data Fig. 10l). Markers of the stem-like HPCS population *SLC4A11*, *HMGA2* and *ITGA2* were significantly positively correlated with *IL34* expression in *KRAS*^G12D^ and *KRAS*^G12V^ tumors, confirming the positive association between stem-like cancer cells and IL-34 observed in the *SftpcCre* mouse model (Fig. 6p, Extended Data Fig. 10m-n). Furthermore, analysis of a dataset of human alveolar epithelial cells treated with IGF1 or vehicle revealed that IGF1 induces the expression of genes that prevent ferroptosis such as *GPX1* or *ACSL3* (Fig. 6q); ferroptosis was the top pathway annotated in WIKIPATHWAYS among genes significantly induced by IGF1 in this dataset (*adj.p*.=1.25×10^-5^), suggesting the antioxidant effect of IGF1 is conserved in human alveolar epithelial cells. Our results have shown that the expression of *IL34* and *IGF1* is associated with poor survival and the maintenance of stem-like cancer cells in human and murine KRAS-mutant lung adenocarcinomas, as well as in other solid tumors and suggest novel therapeutic approaches to treated lung cancer.

Here we have shown that lung tumor-associated, resident macrophages exhibit gene expression signatures of proliferation, oxidative phosphorylation, fatty acid metabolism and immune suppression that confer poor survival on a subset of lung cancer patients. In particular, *KRAS*^G12D^ mutations in mouse lung cells lead to highly penetrant resident macrophage expansion and a mutual IL-34-IGF1-dependent partnership between resident macrophages and lung tumor cells. This partnership has its origins in later embryonic development, as pre-natal ATII cells express a surge of *Il34* in association with the initiation of lung alveolar macrophage expansion. Notably, human *KRAS*^G12D/V^ lung cancers expressing high *IL34* or *IGF1* each exhibit a poor prognosis. IL-34 is a critical self-renewal and growth factor for resident macrophages such as microglia and Langerhans cells^23–25^. Its expression is also upregulated in several different tumor types, where it is associated with poor survival^43–45^. This cytokine maintains resident macrophages in a non-inflammatory state geared toward tissue repair, synaptic pruning, and the maintenance of immune privilege. The upregulation of IL34 by lung tumor cells was unexpected, as GM-CSF, rather than IL-34, is normally required for AM development^29–30^. However, IL-34 and not GM-CSF was clearly associated with selective expansion of AMs in the mouse GEMMs and with TAAM markers in human KRAS-mutant lung cancers.

Our studies have shown that resident macrophages proliferate in KRAS^G12D^ mutant lung tumors under the control of tumor cell-derived IL-34. Resident macrophages then express IGF-1, thereby promoting IGF-1-dependent tumor cell protection from stress responses and ferroptosis, leading to tumor progression. In the autochthonous, genetically engineered *KRAS^G12D^* and *KRAS^G12D^;p53−/−* tumor models studied here, alveolar macrophages are the major population of myeloid cells in lungs, increasing as tumor growth progresses. These results differ somewhat from studies in which tumor growth is initiated by intravenous implantation of *KRAS^G12D^;p53−/−* tumor cells or by installation of adenovirus-expressed Cre in transgenic KRAS^G12D^;p53−/− mouse lungs^10^. In those models, alveolar macrophages do not increase, while bone marrow derived macrophages do increase, during tumor progression^10^. However, other recent lung tumor models have demonstrated increases in AMs that enhance cancer progression^19^. Intratracheally administered lentivirus-Cre in KRAS^G12D^ mice promoted increases in both AMs and bone marrow derived macrophages; therapeutically-mediated reduction in these populations reduced tumor progression^18^. Additionally, mutant EGFR-driven lung adenocarcinoma induces GM-CSF-driven expansion of AMs that enhance tumor growth via cholesterol signaling^19^. These distinctions could be due to difference in microenvironments in the lungs of each model; in autochthonous models used here, mutations are encoded in lung epithelia during embryonic development and their expression is induced in SPC-Cre or CC10Cre expressing cells upon exposure to tamoxifen. In the implanted and virally induced models, introduction of a virus or tumor cells into the lung may simultaneously stimulate host inflammatory responses that are absent from the autochthonous models, leading to the persistent recruitment of bone marrow derived cells. Alternatively, the tumor cells induced in these models may differ in the cytokines they express; differences in these models may identify nuances in lung cancer microenvironments that can ultimately lead to improved therapeutic developments.

Recent studies revealed that oncogenic mutations, including driver mutations, are common in cancer-free tissues; in one study, over 50% of normal lung tissue samples harbored driver somatic mutations in *KRAS*^40^. Whether mutant lung cell clones progress to malignancy depends on clonal competition and the tissue microenvironment. Here we showed that *KRAS*^G12D^ expression in mouse lung cells leads to highly penetrant AM expansion and a mutual IL-34-dependent partnership between AMs and lung tumor cells; In addition, we found that human early-stage *KRAS*^G12D/V^ lung cancers expressing high *IL34* have a poor prognosis. Early *KRAS*^G12D^ mutant clones may benefit from a partnership with AMs; in human early-stage LUAD, clones that co-opt AMs via IL-34 lead to disease with shorter survival. Further studies may identify modifiable factors, such as environmental damage, that increase the likelihood of the IL-34-IGF1 loop induction^46^.

Similarly, in our analysis of the TCGA LUAD dataset, patients with the *KRAS* mutations G12D and G12V showed a significant association of the IL34-IGF1 axis with shorter survival. Taken together, these studies suggest that different oncogenic mutations utilize distinct ways of reprogramming AMs to support tumor development. Mutant-RAS lung cancer cell lines have been shown to be selectively sensitive to IGF1R inhibition^35^; therefore, AM-produced IGF1 may selectively enhance mutant-KRAS lung malignancies. Taken together, these results show that embryonically-derived resident macrophages and bone marrow derived macrophages play discrete roles in the neoplastic setting. Our studies identify novel strategies to control tumor growth by targeting resident macrophage proliferation and interactions with tumor cells.

## Methods

### Animals

All animal studies below were performed with the approval of the Institutional Animal Care and Use Committees and Institutional Biosafety Committees of the University of California, San Diego. Mice were housed in specific pathogen–free conditions on a 12-h light/dark cycle with ad libitum access to food and water. Unless otherwise stated, both sexes were used and randomly assigned to experimental groups. Littermates were used as controls where possible. Health status and body weight were monitored at least weekly and three time a week when tumors were present. 6-8-week-old C57BL/6J (strain #000664 RRID:IMSR_JAX:000664) and Rosa26;LSL-EYFP (*B6.129X1Gt(ROSA)26Sor^tm1(EYFP)Cos^* strain #006148 RRID:IMSR_JAX:020940) animals were purchased from Jackson Laboratories, Bar Harbor, ME and bred at the University of California San Diego. *LSL-Kras^G12D^* (Kras2^tm4Tyj^ RRID:MGI:2429948) and *LSL-p53^R172H+^* (Trp53^tm2.1Tyj^ RRID:MGI:3039264) mice in the C57BL/6 background were maintained in the Varner lab at the University of California, San Diego. *SftpcCreER (B6.129S-Sftpctm1(cre/ERT2)Blh/J* RRID:MGI:5661884), *CC10CreER* (*B6N.129S6(Cg)Scgb1a1^tm1(cre/ERT)Blh/^J*, RRID:IMSR_JAX:016225I) and *LSL-p53^loxp/loxp^* (B6.129P2-*Trp53^tm1Brn^*/J RRID:IMSR_JAX:008462) mice were previously described^12^ and maintained at the Onaitis laboratory at the University of California San Diego. *Il34^LacZ/LacZ^* mice were from Marco Colonna (Washington University) and were crossed with *SftpcCreER and LSL-Kras^G12D^* animals at the University of California San Diego.

### Genetically engineered mouse models of lung adenocarcinoma

*SftpcCreER;Rosa2c-fGFP, SftpcCreER;Rosa26-fGFP;LSL-Kras^G12D^*, *CC10CreER;Rosa26-fGFP, CC10CreER;Rosa26-fGFP;LSL-Kras^G12D^*, *CC10CreER; Rosa2c-fGFP;LSL-Kras^G12D^;LSL-Trp53^loxp/loxp^* and *CC10CreER;LSL-Kras^G12D^;LSL-Trp53^R172H^* genetically engineered mouse models of lung tumor development were obtained by crossing individual strains listed above at the Moores UCSD Cancer Center. To induce tumor development, *SftpcCreER; LSL-Kras^G12D^* mice and control *SftpcCreER mice* were administered tamoxifen (2 mg) in corn oil once at 6 weeks of age. *CC10CreER, CC10CreER;LSL-Kras^G12D^* and *CC10CreER;LSL-Kras^G12D^;LSL-Trp53^R172H^* mice were administered 2 mg tamoxifen (#T5648 Sigma) in corn oil every other day for eight days beginning at 6 weeks of age. To evaluate tumor development and inflammation over time, *SftpcCreER; Rosa2c-fGFP;LSL-Kras^G12D^* animals were euthanized and lungs were isolated for further study at 2-or 5-weeks post tamoxifen administration. *CC10CreER; Rosa2c-fGFP;LSL-Kras^G12D^* and *CC10CreER; Rosa26-fGFP;LSL-Kras^G12D^;LSL-Trp53 ^loxp/loxp^* were euthanized and lungs were isolated for further study at regular intervals between 4- and 18-weeks post tamoxifen administration. For scRNA sequencing, lungs from *SftpcCreER, SftpcCreER;LSL-Kras^G12D^* and *SftpcCreER;LSL-KrasG12D;IL34*−/− animals were isolated at ages 8-10 weeks of age, and *CC10CreER and CC10CreER;LSL-Kras^G12D^;LSL-Trp53^R172H^* lungs were isolated 13 weeks post-tamoxifen injection. In some experiments, *CC10CreER;LSL-Kras^G12D^;LSL-Trp53^R172H^* or *SftpcCreER;LSL-Kras^G12D^* animals were treated with 200μg neutralizing rat anti-mouse IL34 (IgG2A clone #780310 R&D Systems MAB5195 RRID:AB_3094866), IgG2A control (BE0089-R001) or saline 2x per week for 2-3 weeks.

### Bronchoalveolar lavage

To collect cells and fluid from the bronchoalveolar spaces of *SftpcCreER* and *SftpcCreER; LSL-Kras^G12D^* mouse lungs, mice were euthanized and 1 ml of PBS was injected into the lungs by intratracheal cannulation and then aspirated into a 15 ml conical tube. The resulting lavage was centrifuged 7 min at 300×g at 4°C. Supernatants (bronchoalveolar lavage fluid, BALF) was removed and centrifuged at 10,000 rpm at 4°C to remove residual cells for further cytokine measurement by ELISA assays, as described below. Cells from lavages were resuspended in 1 ml PBS and re-centrifuged twice. A small sample was centrifuged using a Cytospin centrifuge (ThermoFisher) for morphological analysis by Wright Giemsa staining and counting. Total RNA was extracted from the remaining cells from three biological replicates for each condition using the Qiagen RNeasy kit according to the manufacturer’s instructions. Messenger RNA purification, fragmentation, construction of sequencing libraries and sequencing were performed using the Illumina Pipeline.

### Lung/tumor dissociation and preparation of single-cell suspensions

Lungs with and without tumors were excised and immediately placed in a sterile Petri dish on ice. Using sterile scissors, the tumors were cut into small fragments (approximately 4 mm^2^ in size) and transferred into Miltenyi gentleMACS™ C Tubes (# 130-093-237). Tumor dissociation was performed using the Miltenyi Tumor Dissociation Kit (#130-096-730) on a Miltenyi gentleMACS™ Octo Dissociator with Heaters (RRID:SCR_020271), with the appropriate pre-set tumor dissociation program. Following enzymatic and mechanical dissociation, cell suspensions were passed through a 70 μm cell strainer into a 50 mL conical tube to remove debris and large aggregates. Red blood cells were lysed by adding 1× red blood cell lysis buffer (Pharm Lyse, #555899, BD Biosciences) and incubating for the manufacturer-recommended time at room temperature. The suspension was then passed through a second 40 μm cell strainer to ensure a uniform single-cell suspension. Cells were washed once with PBS containing 2% FBS, centrifuged at 300 × g for 5 minutes at 4°C, and resuspended in PBS containing 2% FBS for counting. Cell counts were performed using the Revvity Cellometer Ascend cell counter. The resulting single-cell suspensions were immediately used for downstream flow cytometry analysis or bulk and single cell sequencing.

### Immunohistochemistry

Formalin-fixed, paraffin-embedded (FFPE) lung tissue sections (5 μm thickness) were mounted on charged glass slides and dried overnight at 37°C. Sections were deparaffinized in xylene (2 × 5 min) and rehydrated through a graded ethanol series (100%, 95%, 70%; 2 min each) followed by rinsing in distilled water. Antigen retrieval was performed by heating slides in Diva Decloaker (Biocare Medical DV2004MX) under steam for 30 minutes at 95°C. Slides were cooled to room temperature in the retrieval buffer and rinsed in PBS-T (PBS containing 0.05% Tween-20). Endogenous peroxidase activity was quenched by incubating sections with Bloxall peroxidase blocking solution (Vector Laboratories SP-6000-100) for 10 min at room temperature, followed by two washes in PBS-T. To minimize nonspecific antibody binding, sections were blocked with 2.5% normal goat serum for 20 min at room temperature in a humidified chamber. Sections were then incubated with 1:250 rat Anti-F4/80 (eBioscience #14-4801-85, clone BM8, RRID:AB_467559) or anti-SiglecF (ThermoFisher #PA5-11675, RRID: AB_2189281) in 2.5% normal goat serum in Phosphate buffered saline (PBS) for 1h at room temperature or overnight at 4°C. After three washes in PBS-T, sections were incubated with goat anti-Rat IgG ImmPRESS HRP Polymer (Mouse Adsorbed, Vector Laboratories MP-7444-15) or ImmPressHRP Horse anti-Rabbit Ig Polymer (Vector Laboratories MP-7401) for 30 min at room temperature. After PBS-T washes, antibody binding was visualized using the DAB Peroxidase Substrate Kit (Vector Laboratories SK-4105) according to the manufacturer’s protocol, monitoring under a microscope for optimal signal development. The reaction was stopped by immersion in distilled water. Sections were counterstained with hematoxylin (Vector Laboratories H-3404) for 1 min, rinsed in running tap water, blued in 0.1% sodium bicarbonate, dehydrated through graded ethanol series, cleared in xylene, and mounted with Cytoseal permanent mounting medium. Negative controls were prepared by omitting the primary antibody. All slides were examined and imaged using a brightfield microscope under identical exposure settings. Slides were scanned a Leica AT2 Aperio scanner (RRID:SCR_021256). Image analysis and quantification of cells/biomarkers was performed using Qpath open-source digital pathology analysis software^48^ and Image J open-source imaging software.

Two-color fluorescent immunohistochemistry was performed to detect Ki67 and F4/80 or MERTK and F4/80 in the same tissue. After processing of tissues as described above with either rabbit anti-mouse anti-Ki67 from Abcam (ab15580, RRID:AB_443209) or rabbit anti-mouse MERTK from Thermo Fisher (clone RM469, MA5-44594, RRID:AB_2926724), tissues were incubated with ImmPressHRP Horse anti-Rabbit Ig Polymer (Vector Laboratories MP-7401), washed in TBST and incubated in AlexaFluor488 Tyramide reagent (Vector Laboratories B40953) for 10 min. Slides were then incubated in 1:200 rat anti-F4/80 (eBioscience #14-4801-85, clone BM8, RRID:AB_467559), ImmPress HRP Goat anti-Rat IgG Polymer Detection Kit (MP-7444-15) and AlexaFluor 594 were viewed on a Nikon TE2000 epifluorescence microscope (RRID:SCR_023161) and scanned on a Akoya Phenoimager RRID:SCR_0237720). Image analysis and quantification of cells/biomarkers was performed using Qpath open-source digital pathology software^48^.

### Flow cytometry staining and analysis

Single-cell suspensions were prepared at a concentration of 1 × 10^6^ cells in 100 μL total volume of PBS containing 2mM EDTA and 2.5% Bovine Serum Albumin (BSA). Cells were incubated with LIVE/DEAD™ Fixable Blue Dead Cell Stain (Invitrogen # L34962) and FcR Blocking Reagent (BD Pharmingen #553142) at room temperature for 15 minutes to exclude non-viable cells and block Fc receptor–mediated nonspecific binding. After blocking, cells were stained with fluorescently conjugated primary antibodies against cell surface markers, incubating for 30 minutes at 4°C in the dark. For intracellular staining, cells were fixed, permeabilized using Transcription Factor Staining Buffer Set (#00-5523-00, ThermoFisher) and then incubated with fluorescently labeled antibodies. Primary antibodies used for cell surface marker staining were: CD45-AF700 (1μg/mL, clone 30F11, #56-0451-82, Invitrogen RRID: AB_891454), anti-CD11b-AF288 (M1/70, Biolegend #101219, AB_830641 or PerCP-Cy5.5 or Biolegend #101227, RRID: AB_893233), F4/80-SB670 (5μg/mL, clone A3-1, #MCA4975SB670, BioRad RRID: AB_3101056), or F4/80-BV650 (5 μg/mL, clone BM8, #416-4801-82 RRID:AB_2925697), Ly6C-RB705 (1μg/mL, clone AL-21, # 570264, Becton Dickinson), Ly6C-SB670 (1ug/ml, clone AL-21, #570264, Becton Dickinson, RRID:AB_3685625) or Ly6C-BV421 (1ug/ml, clone AL-21, #562727, Becton Dickinson, RRID:AB_2737748), Ly6G-PE-Cy7 (1μg/mL, clone 1A8, #560601, Becton Dickinson RRID:AB_1727562), CD11c-APC (1μg/mL, clone HL3, #550261, Becton Dickinson, RRID: AB_10562405), CD206-AF594 (1ug/ml, clone C068C2, #141726, Biolegend, RRID:AB_2563300), SiglecF-BV785 (1ug/ml clone E50-2440 #740956 Becton Dickinson, RRID:AB_2740581), EpCAM-BV786 (clone G8.8, #118245 Biolegend RRID: AB_2860639), MHCII-eF450 (clone AF6-120.1, #48-5320-82, ThermoFischer, RRID:AB_10669941) and T1a-PE (8.1.1, #127407 Biolegend, RRID: AB_2161929) for 30 min at RT in the dark. The cells were washed twice in PBS containing 2% FBS then were sorted into alveolar macrophages (live cells, CD45+autofluorescent+F4/80+Ly6C-Ly6G-CD11c+) and alveolar epithelial cells (live cells, CD45-EpCAM^int^ MHCII+T1a-). After staining, cells were washed twice with cold PBS containing 2% FBS, centrifuging at 300 × g for 5 minutes at 4°C between washes. The cell pellet was then resuspended in Stabilizing Fixative (#338036, Becton Dickinson) for preservation. Multicolor FACS Analysis was performed on BD LSR Fortessa 18-color analyzer (Sanford Consortium for Regenerative Medicine) or on the 11 color BD Facs Aria at the Moores Cancer Center Flow Cytometry core. Compensation was applied using single-stained controls, and fluorescence minus one controls were included as appropriate. All data analysis was performed using the flow cytometry analysis program FlowJo (Becton Dickinson/Treestar; RRID:SCR_008520).

### Ferroptosis assessment

To assess ferroptosis/lipid peroxidation in tumor cells, dissociated tumor cells were stained with anti-CD45 Alexa700 (clone 30-F11, Invitrogen #56-0451-82, RRID:AB_891454), anti-EpCAM+ APC (clone G8.8, Biolegend #118213, RRID:AB_1134105) and BODIPY 581/591 C11 Lipid Peroxidation Sensor (Cell Signaling Technology #95978). Multicolor FACS Analysis was performed on the 11 color BD Facs Aria at the Moores Cancer Center Flow Cytometry core. Lipid peroxidation in CD45⁻EpCAM⁺ tumor cells was quantified as the ratio of oxidized to reduced BODIPY C11 fluorescence, measured as MFI in the FITC/green channel (510-530 nM emission) relative to the PE/red-orange channel (575-590 nM emission).

### Flow Cytometric Analysis of CSF1R and CSF2R Expression

Single-cell suspensions of 8-week-old *SfptcCre* and *SfptcCreKras*^G12D^ lungs were first stained with Live/dead™ Fixable Aqua Dead Cell Stain Kit (Thermo Scientific,# L34966) to exclude dead cells. Following viability staining, cells were incubated with 1 ug/ml fluorochrome-conjugated antibodies against CD45 (clone 30-F11, AF700, Invitrogen, #56-0451-82, RRID: AB_891454), CD11b (clone M1/70, PerCP-Cy5.5. Biolegend, #101227, RRID: AB_893233), Ly6G (clone 1A8, PECy7, Becton Dickinson Biosciences, #560601, RRID: AB_1727562), Ly6C (clone AL-21, BV421, Becton Dickinson Biosciences, #562727, RRID: AB_2737748), F4/80 (clone C1:A3-1, BV670, Bio-Rad, #MCA497SBV670, RRID: AB_3101056), SiglecF (clone E50-2440, BV786, Becton Dickinson, #740956, RRID: AB_2740581) and CD11c (clone N418, APC/Cy7, Biolegend <u>#117323</u> RRID: <u>AB_830646</u>) to identify the AMs populations. To assess receptor expression, cells were additionally stained with antibodies against CSF1R/CD115 (clone AFS98, APC, Biolegend, #135509, RRID: AB_2085222) or CSF2R/CD131 (clone REA193, APC, Miltenyi Biotec, #130-103-076, RRID: AB_2654863). Following antibody staining, cells were washed twice with staining buffer and resuspended for acquisition on a flow cytometer. Compensation controls and fluorescence-minus-one (FMO) controls were used where appropriate. Data were analyzed using FlowJo software (BD Biosciences). AMs populations were identified using a sequential gating strategy. Expression of CSF1R (CD115) and CSF2R (CD131) was quantified as the percentage of positive cells and mean fluorescence intensity (MFI) within the indicated populations.

### Isolation of macrophages and epithelial cells from *SftpcCre;KrasG12D* animals

Lungs were dissociated into single cells as described above. Dissociated cells from lungs were centrifuged at 450xg for 4 min at 4°C, incubated with ACK buffer (A1049201, Gibco) to lyse red blood cells and 1 min at room temperature, washed with 1x PBS, and counted. Resulting cell suspensions were incubated for 10 min at RT in the dark with eBioscience™ Fixable Viability Dye eFluor™ 780 (1:1000, #65-0865-14, eBioscience) diluted in 1x PBS, in the presence of Fc block (#553141, BD) and true-stain monocyte blocker (#426102, Biolegend). After washing with PBS containing 2% FBS, cells were stained with the following antibodies: CD45-AF700 (1μg/mL, clone 30-F11, #56-0451-82, Invitrogen RRID: AB_891454), F4/80-SB670 (5μg/mL, clone A3-1, #MCA4975SB670, BioRad RRID: AB_3101056), Ly6C-RB705 (1μg/mL, clone AL-21, # 570265, Becton Dickinson, RRID: AB_3685625, Ly6G-PE-Cy7 (1μg/mL, clone 1A8, # 560601, BD RRID:AB_1727562), CD11c-APC (1μg/mL, clone HL3, # 550261, BD RRID: AB_10562405), EpCAM-BV786 (clone G8.8, #118245 Biolegend RRID: AB_2860639),MHCII-eF450 (clone AF6-120.1, #48-5320-82, ThermoFischer, RRID:AB_10669941) and T1a-PE (8.1.1, #127407 Biolegend, RRID: AB_2161929) for 30 min at RT in the dark. The cells were washed twice in PBS containing 2% FBS then were sorted into alveolar macrophages (live cells, CD45+autofluorescent+F4/80+Ly6C-Ly6G-CD11c+) and alveolar epithelial cells (live cells, CD45-EpCAM^int^ MHCII+T1a-) on a FACSymphony A6 sorter (Becton Dickinson), or at the Moores Cancer Center on a BD FACSAria II SORP (High speed, 13-color, 4-laser cell Sorter), in the Moores Cancer Center flow cytometry shared resource. Cells were used for bulk RNA sequencing or cultured ex vivo.

### In vitro culture of sorted alveolar macrophages

Alveolar macrophages sorted from 3 separate *SftpcCreERT2;Kras^G12D^* mice were cultured ex vivo in DMEM containing 20% fetal bovine serum and 50 ng/ml recombinant mIL34 (5195-ML-010, R&D Systems) in the presence or absence of 10 µg/ml anti-mIL34 (MAB5195, Clone #780310, R&D Systems RRID:AB_3094866) for 7 days. Cells monolayers were washed with PBS and total RNA was extracted using miRNeasy Micro Kit (Qiagen #217084) according to manufacturer’s instructions. RNA samples with RIN over 7.0 were subjected to library generation and high-throughput sequencing.

### Cytokine-Induced Alveolar Macrophage Proliferation Assay

Sorted alveolar macrophages were cultured in complete RPMI-1640 medium supplemented with 10% fetal bovine serum and 1% penicillin-streptomycin. Cells were seeded at a density of 3,000 cells per well in 96-well tissue culture plates and stimulated with recombinant mouse IL-34 (R&D Systems, #5195-ML-010) at 50, 100, or 200 ng/mL; recombinant mouse M-CSF (PeproTech, #315-02) at 25, 50, or 100 ng/mL; or recombinant mouse GM-CSF (PeproTech, #315-03) at 25, 50, or 100 ng/mL. Untreated cells cultured in complete medium served as controls. Cell proliferation was assessed at 0, 24, 48, and 72 hours after cytokine stimulation. At each time point, cell growth was quantified using a CCK8 assay, according to the manufacturer’s instructions.

Following collection of culture supernatants for ELISA analysis, 100 μL of fresh complete culture medium containing Cell Counting Kit-8 (CCK-8) reagent was added to each well according to the manufacturer’s instructions. Cells were incubated at 37°C for 2 h, and absorbance was measured at 450 nm using a microplate reader. Cell viability and proliferation were expressed as relative absorbance values normalized to the untreated control group. The survival ratio for each treatment condition was calculated as the percentage of viable cells relative to vehicle-treated controls

### IGF1 Pathway Analysis

Sorted tumor cells were seeded at a density of 3,000 cells per well in 96-well tissue culture plates and allowed to recover overnight before further treatment. To evaluate the role of IGF1 signaling in tumor cell proliferation and IL-34 secretion, cells were treated with the following conditions: Vehicle control (0.1%DMSO for inhibitors or saline for antibodies), 100 ng/mL recombinant mouse IGF1 (ThermoFisher PMG0075), anti-IGF1 neutralizing antibody (10 μg/mL R&D Systems #MAB791, RRID:AB_2122252), or 1 µM Linsitinib (OSI-906, Selleckchem #S1091) or NVP-AEW541 (Selleckchem #S1034) in 0.1% DMSO. For IGF1 stimulation experiments, recombinant mouse IGF1 was added to the culture medium at the indicated concentration and incubated for 48h. For inhibition studies, anti-IGF1 antibody or IGF1R inhibitors were added at the time of cell seeding and maintained throughout the treatment period. Cells were cultured under standard conditions (37°C, 5% CO₂) for 48 h.

### Cell Proliferation Assay

Following collection of culture supernatants for ELISA analysis, 100 μL of fresh complete culture medium containing Cell Counting Kit-8 (CCK-8) reagent was added to each well according to the manufacturer’s instructions. Cells were incubated at 37°C for 2 h, and absorbance was measured at 450 nm using a microplate reader. Cell viability and proliferation were expressed as relative absorbance values normalized to the untreated control group. The survival ratio for each treatment condition was calculated as the percentage of viable cells relative to vehicle-treated controls.

### Tumor Cell–Alveolar Macrophage Co-culture

To assess AM proliferation in the presence of tumor cells, sorted tumor cells were seeded into the upper inserts of a 24 Transwell plate, while sorted alveolar macrophages (AMs) were seeded into lower chambers. To assess tumor cell proliferation in the presence of alveolar macrophages, alveolar macrophages were seeded into the upper inserts of a 24 Transwell plate, while sorted tumor cells were seeded into lower chambers. Both cell populations were cultured in complete RPMI-1640 medium supplemented with 10% Fetal Bovine Serum and 1% penicillin-streptomycin and allowed to adhere overnight prior to treatment. Tumor cell (upper)-alveolar macrophage (lower) cultures were incubated in the presence of medium or anti-IL-34 neutralizing antibodies (10 μg/mL neutralizing rat anti-mouse IL34 (IgG2A clone #780310 R&D Systems MAB5195 RRID:AB_3094866). Alveolar macrophage (upper)-tumor cell (lower) cultures were incubated in the presence of vehicle (0.1% DMSO), 10 µg/ml recombinant mouse anti-IGF1 neutralizing antibody (R&D Systems #AF791, RRID:AB_2248752), OSI-906, Selleckchem #S1091) or NVP-AEW541 (Selleckchem #S1034). Cells were cultured in a humidified incubator at 37°C with 5% CO₂ for 48 h.

### Collection of Conditioned Medium and IGF1 ELISA

After 48 h of co-culture, Transwell inserts containing cells were carefully removed. Culture media from both the upper and lower chambers were pooled and mixed thoroughly to generate a combined conditioned medium sample representing secreted factors from both tumor cells and alveolar macrophages. A total of 500 μL of pooled conditioned medium was collected from each well and centrifuged briefly to remove cellular debris. Supernatants were stored at −80°C until analysis. IGF1 concentrations were quantified using a Mouse IGF1 Quantikine ELISA Kit (R&D Systems, #MG100, RRID: AB_2827989) according to the manufacturer’s instructions. IL-34 concentrations were quantified using a Mouse IL-34 Quantikine ELISA Kit (R&D Systems #M3400, RRID:AB_2820306) Briefly, standards and samples were added to pre-coated ELISA plates and incubated as recommended by the manufacturer. Following washing steps, detection antibody, enzyme conjugate, and substrate solution were sequentially applied. Absorbance was measured at 450 nm with wavelength correction at 570 nm using a microplate reader. Cytokine concentrations were calculated using a standard curve generated from recombinant protein standards.

### Bone marrow derived macrophage differentiation and culture

Bone marrow derived cells (BMDC) were aseptically harvested from 8-week-old C57/BL6 female mice by flushing leg bones of euthanized mice with phosphate buffered saline (PBS), 0.5% BSA, 2mM EDTA, incubating in red cell lysis buffer (155 mM NH_4_Cl, 10 mM NaHCO_3_ and 0.1 mM EDTA) and centrifuging over Histopaque 1083 (Sigma-Aldrich #10831) to purify the mononuclear cells. Approximately 5×10^7^ BMDC were purified by gradient centrifugation from the femurs and tibias of a single mouse. Purified mononuclear cells were cultured in DMEM + 20% serum + 50 ng/ml M-CSF (PeproTech #315-02 RRID:AB2744479). Day 7 macrophages were serum starved overnight by culturing in DMEM with 0.5% FBS and stimulated for 0, 1, 2, 3, 5 10, 15, 30 min 50ng/ml M-CSF or for 0, 2, 5, 10. 30 min IL-34, then were solubilized on ice with RIPA buffer containing protease and phosphatase inhibitors.

### Immunoblotting

Lysates from bone-marrow-derived macrophages were electrophoresed on 8% Bis-tris minigels and transferred to PVDF membranes using an iBlot Western transfer system (ThermoFisher). After blocking non-specific protein binding sites by incubation for 2 hours room temperature in 5% bovine serum albumin (BSA) in Tris buffered saline containing 0.1% Tween 20, pH 7.4 (TBST). Blots were then incubated with gentle agitation at 4°C overnight in the presence of primary antibodies diluted 1:1000 in 5% BSA in TBST. Antibodies were rabbit anti-phosphoTyr723 CSF1R (clone 49C10, #3155, RRID:AB_2085229), rabbit anti-CSF1R (clone D3O9X, 67455, RRID:AB_2799725), rabbit anti-phosphoSer473 Akt (clone D9E, #4060, RRID:AB_2315049), Akt (clone C67E7, #4691, RRID:AB_915783), phosphoThr202/Tyr204 Erk (clone 197G2, #4377, RRID:AB_331775), and Erk (clone 137F5, #4695, RRID:AB_390779) from Cell Signaling Technologies. Membranes were washed three times for 10 min each with TBST, then incubated with Horse Radish Peroxidase-conjugated goat anti-rabbit IgG (heavy plus light chain) antibodies (Cell Signaling Technology #7074, RRID:AB_2099233) at 1:5000 in TBST+ 5% BSA for 1 hour at room temperature. Blots were washed three times again for 10 min each with TBST prior to detection with a 1:1 mixture of peroxide and luminol prior to imaging on an iBright Cl1500 chemiluminescence imaging system (Invitrogen, RRID:SCR_026565).

### Cytokine quantification

The protein concentrations in lysates of lungs from *SftpcCreERT2;Kras^G12D^* and *CC10CreERT2;LSL-Kras^G12D^;LSL-Trp53^R172H^* mice and cell culture supernatants from in vitro cultured Lewis lung carcinoma cells (LLC) were determined using a BCA Protein Assay (ThermoFisher #A55865) according to manufacturer’s instructions. Tumor cell supernatants (100 µL) or 500 µg total protein from tumors were used in ELISAs to detect cytokines. Cytokine concentrations in samples were determined using R&D Systems Quantikine ELISA kits for Mouse IL-34 (R&D Systems #M3400, RRID:AB_2820306), Mouse/Rat IGF-I/IGF-1 (R&D Systems, #MG100, RRID: AB_2827989), Mouse M-CSF (R&D Systems #MMC00B, RRID:AB_2894179), and Mouse GM-CSF (R&D Systems #MGM00, RRID:SCR_007050), following the manufacturer’s instructions. Protein expression was normalized to total volume (of supernatants) or 100 µg total protein (for tumor lysates).

### Bulk RNA sequencing of sorted cells from genetically engineered lung cancer models

Total RNA was extracted from mouse lungs using miRNeasy Micro Kit (Qiagen #217084) according to manufacturer’s instructions. RNA samples with RIN over 7.0 were subjected to library generation and high-throughput sequencing. Raw sequencing reads (~ 40 M per sample) were first processed for adapter trimming using trim-galore (version 0.6.10). The trimmed reads were then aligned to the mouse reference genome (mm39) using STAR aligner (version 2.7.11b). Gene counts were obtained using FeatureCounts (Subread, version 2.0.6). To identify differentially expressed genes, differential gene expression analysis was conducted using DESeq2 (version 1.48.1). Genes with an adjusted p-value < 0.05 were considered significantly differentially expressed. For gene set enrichment analysis (GSEA), ranked gene lists were generated based on log2 fold changes, enrichment scores were calculated using clusterProfiler (version 4.16.0).

### 10X Genomics single-cell RNA-sequencing library preparation

Isolated single-cell suspensions from LLC tumors, *SftpcCreER, SftpcCreER;Il34*−/−*, CC10CreER, CC10CreER;Kras^G12D;^p53^R172H^, SftpcCreER;Kras^G12D^, SftpcCreER;KrasG12D;Il34*−/− were resuspended in DMEM at a concentration of about 750 cells/µl. Cells were counted with a Cellometer Ascend Automated Cell Counter (Revvity) to determine their concentration. Single-cell RNA-sequencing libraries were prepared using Chromium Connect with the Next GEM Automated Single Cell 3′ reagent kit v3.1 (10x Genomics RRID:SCR_025146), following the manufacturer’s protocol. Up to 24,000 cells were loaded onto the Next GEM Chip G. The size of the sequencing libraries was measured using the 4200 TapeStation system (Agilent, RRID:SCR_018435), and the concentration of each individual library was quantified by Qubit dsDNA High Sensitivity assay kit (Thermo Fisher Scientific). After preparation, final libraries were sequenced on the NovaSeq 6000 (RRID:SCR_016387) or NovaSeq X Plus System platform (Illumina, RRID:SCR_024568).

### Single-cell RNA-sequencing analysis

Sequencing data pre-processing of the FASTQ files were performed using Cell Ranger (version 9.0.1). Raw reads were aligned to the mouse (GRCm39) 2024-A reference. Samples were then preprocessed using Seurat 5.1.0 (RRID:SCR_016341) workflow, including NormalizeData, ScaleData, FindVariableFeatures, RunPCA, FindNeighbors, RunUMAP, FindClusters; quality control was performed using Seurat.^49^ Batch-correction was performed with Harmony (version 1.2.3)^50^ for integration of *CC10Cre, CC10Cre;Kras^G12D^p53^fl/fl^* (isolated 13 weeks after tamoxifen induction) and *SftpcCreER* and *SftpcCreER;Kras^G12D^* data sets.

The Seurat implementation of UMAP using the package UWOT (Melville J (2024) uwot: The Uniform Manifold Approximation and Projection (UMAP) Method for Dimensionality Reduction. R package version 0.2.2, https://CRAN.R-project.org/package=uwot) was used for dimensionality reduction with default parameters. For differential analysis of major cell type clusters, pseudobulk analysis using DESeq2 1.44.0 was used with a single-factor two-group design with 2-4 replicates per group.^51^ Plots were generated using Seurat functions DotPlot, ViolinPlot, DimPlot, and FeaturePlot. Heatmaps were plotted using the ComplexHeatmap package^52^. To perform gene set enrichment analysis of single-cell RNA sequencing clusters, pseudobulk DESeq2 (RRID:SCR_015687) output was used as input for the GSEA (RRID:SCR_003199) program using the pre-ranked mode^53,54^. Default settings in the pre-ranked mode were used. The ggplot2 (RRID:SCR_014601) function geom_point was used for plotting GSEA outputs. MSigDB version 2024.1 was used^54^.

Datasets from single experiments with matched controls (such as *CC10CreER;Kras^G12D;^p53^fl/fl^* and *CC10Cre* controls) were analyzed by processing individual samples using Seurat then merging the samples into a combined Seurat object. Normalization scaling, dimensionality reduction, clustering and downstream analyses were performed using the merged Seurat object. Datasets from independent experiments were processed by merging or integrating the individual experiment’s Seurat objects. Specifically, the following datasets of their combinations were analyzed by merging: *CC10CreER and CC10CreER;Kras^G12D^;p53^fl/fl^* (Figure 2a), *SftpcCreER*, *SftpcCreER;Kras^G12D^* (Figure 2n), *SftpcCreER*, *SftpcCreER;Kras^G12D^*, *SftpcCreER;Il34*^−/−^, *SftpcCreER*; *Kras^G12D^*;*Il34*^−/−^ (Figure 4k), *Map17CreER* (GSE156232, Extended Data Figure 4), *CC10CreER;Kras^G12D^;p53^fl/fl^* and *SftpcCreER;Kras^G12D^* anti-IL34 and control treated samples (Extended Data Figures 8–9). The following datasets were analyzed by integration using the experiment of origin as the integration variable: integration of *Map17CreER* and *CC10CreER* experiments (Extended Data Figure 4). The following dataset was integrated using patient of origin as the integration variable: Leader et al. human lung cancer samples (Figure 6).^17^ Integration methods were selected for each integration based on considerations of computational feasibility and integration performance. Integration performance was evaluated by inspecting successful integration of major cell types into major cell type clusters (such as T cells, fibroblasts, endothelial cells) and preservation of some degree of sample-wise subcluster heterogeneity. Multiple methods for which Seurat implementations are available were considered: CCA, RPCA, SCTransform v2, Harmony, Sketch integration using RPCA. The Leader et al. dataset was integrated using the sketch + RPCA integration method.

### Identification of tumor-associated alveolar macrophage marker genes

To obtain a list of mouse tumor-associated alveolar macrophage marker genes, genes upregulated in tumor-enriched vs normal tissue-enriched alveolar macrophage clusters were identified in the *CC10CreER*/*CC10CreER;Kras^G12D^;p53^fl/fl^* and *SftpcCreER/SftpcCreER;Kras^G12D^* datasets by comparing Cluster 0 to Cluster 1 and Cluster 0 to Cluster 10, respectively. The genes were filtered by adjusted p-value (*p*<0.05), fold change (log_2_(fold change)>0 in tumor-enriched cluster) and minimum expression of 25% in the tumor-enriched cluster in both datasets. Finally, a list of 1009 consensus genes upregulated in the tumor-enriched AM cluster in both datasets was obtained and used for TAAM marker gene expression assessment. To establish the dependence of these TAAM markers on IL-34, the *SftpcCreER;KrasG12D;Il34^−/−^* dataset and the associated *Il34*^+/+^ controls were utilized (Figure 4k,n). Differential expression analysis identified 1441 genes significantly downregulated in *Il34*^−/−^ enriched TAAM cluster (Cluster 2) and *Il34*^+/+^ enriched TAAM cluster (Cluster 1) and the overlap between these genes and TAAM markers was determined and presented in Extended Data 8e.

### Analyses of intercellular signaling in scRNA-seq data (NicheNet and CellChat)

To analyze intercellular communication, CellChat v2 (RRID:SCR_021946) was applied to scRNA-seq data using the recommended analysis workflow^31^ using the following functions: netAnalysis_signalingRole_scatter, netVisual_heatmap, netVisual_diffInteraction, netAnalysis_signalingRole_heatmap, netAnalysis_signalingChanges_scatter, netVisual_bubble. R packages Seurat and ggplot2 (RRID:SCR_014601) were used for additional visualization of results. The list of interactions belonging to each pathway as defined by CellChat2 can be accessed through the CellChat R package or online database (http://www.cellchat.org/cellchatdb/). Additional intercellular communication analysis was performed using nichenetr, the R implementation of NicheNet^22^ (available through GitHub as saeyslab/nichenetr). This method was applied to analyze signaling in the *CC10CreER and CC10CreER;Kras^G12D^;p53^fl/fl^* datasets and the nichenetr vignette was followed with the following application details: the reference condition was set as *CC10CreER* (tumor-free lung) and the condition of interest as *CC10CreER;Kras^G12D^;p53^fl/fl^*; the signaling cut-off for prepare_ligand_target_visualization was 0.25; for the analysis in Figure 2g, Cluster 0 was set as the target receiver cluster and all other clusters were considered as potential sender clusters; threshold for expressed genes in get_expressed_genes was set at 0.05.

### Cell type prediction in scRNA-seq data

To assist with mouse cell type identification using marker genes, an automated cell type prediction method was utilized using the ImmGen large-scale transcriptomic dataset and the R packaged celldex and SingleR^55^. Both packages are available through Bioconductor. The celldex package was used to obtain ImmGen reference data using the fetchReference command and specifying “immgen” and “2024-02-26” as parameters. The Seurat object of interest was converted to a singleCellExperiment using the as.SingleCellExperiment function and cell type prediction performed using the SingleR function using label.fine as a parameter i.e. fine labels from the ImmGen dataset. The resulting predictions were plotted using the ComplexHeatmap package^52^ by calculating the most abundant cell type predictions per cluster and plotting the cluster share of each cell type prediction (with 1 being the maximum for each cluster-cell type prediction).

### RNA velocity and trajectory analysis

RNA velocity analysis was performed using velocyto, velocyto.R, scvelo and cellrank packages. Quantification of spliced and unspliced reads was performed using the velocyto^20^ run10x command on the cellranger output BAM file. To apply RNA velocity estimations to existing single cell datasets, analyses were performed for all cells as well as individual sample groups. In both cases cells with matching barcodes in a Seurat object and the velocyto output loom file were identified and the spliced, unspliced and ambiguous reads inserted into the Seurat object. The resulting Seurat object was subjected to analysis by the RunVelocity function from velocyto.R^20^ exported in the HDF5 format using the SaveH5Seurat function from the R package SeuratDisk version 0.0.0.9 (available through Github as mojaveazure/seurat-disk) and converted to the h5ad format (HDF5 AnnData object). The resulting AnnData object was subjected to standard scvelo analysis. Briefly, scvelo functions scv.pp.filter_and_normalize, scv.pp.moments, scv.tl.velocity and scv.tl.velocity_graph were applied sequentially and the velocity vectors plotted using the scv.pl.velocity_embedding_stream function. RNA velocity and expression levels of individual genes were plotted using the scv.pl.velocity function. Analysis of predicted origin and target states was performed using CellRank version 2.0 following the recommended workflow.^21^ For each hypothetical starting cluster as origin, the likely target states were visualized using the vk.plot_random_walks function. Initial and terminal states were predicted using the following functions: cr.estimators.GPCCA, fit, plot_macrostates, predict_terminal_states, predict_initial_states and plot_macrostates.

#### Analysis of IL-34-induced genes in human cells

To obtain a list of genes induced by IL-34 in human monocytes/macrophages, the publicly available bulk RNA-seq dataset GSE151194^47^ was re-analyzed from raw counts available on GEO using DESeq2. All sample groups were included in the DESeq model construction and variance analysis. For differential expression analysis, to obtain IL-34 induced genes, the IL-34 treated samples were compared to the M0 (untreated monocytes) samples. To compare the IL-34-induced transcriptome from monocyte-derived macrophages treated with M1 polarizing conditions, the IL-34 treated group was compared to the M1 (IFNγ and GM-CSF treated) group. Adjusted p<0.05 was used as the significance threshold. R version 4.5 and DESeq2 version 1.48 were used.^51^

#### Analysis of lineage tracing (Map17CreER-CC10creER integration)

To determine whether CC10CreER tumor-associated macrophages phenotypically resemble tissue-resident or bone marrow-derived macrophages, a lineage-tracing dataset of empirically identified bone marrow-derived (Map17-labeled) and tissue-resident (Map17-unlabeled) macrophages was used (GSE156232)^10^. The raw counts matrix files available on GEO were analyzed by Seurat version 5 using a standard single-dataset workflow. The resulting Seurat object was subjected to integration using the sketch RPCA integration approach^56^. Dataset of origin was used as the integration variable i.e. the Map17CreER dataset was integrated with the CC10creER dataset. Following integration, the full dataset was projected onto the sketch-integrated dataset and subjected to downstream analysis using standard workflow.

#### Analysis of human lung tumor scRNA-seq data

To analyze alveolar macrophage transcriptome in human lung cancer at single-cell resolution, the publicly available dataset by Leader et al.^17^ was subjected to re-analysis (GSE154826). Individual patient transcriptome datasets were subjected to standard Seurat processing and cell type annotation and then integrated by patient of origin as the integration variable using the sketch integration approach. This required a 512 GB memory computing node for completion. Metadata available on GEO were added to the Seurat object and alveolar macrophages subjected to downstream analysis.

#### Analysis of LungMAP developing mouse lung dataset

To investigate alveolar type 2 (AT2) cell and alveolar macrophage (AM) transcriptomes during mouse lung development at single-cell resolution, the publicly available LungMAP dataset (LMEX0000004397) was downloaded from the LungMAP Consortium portal (https://www.lungmap.net) (PMID: 37516747). This dataset comprises single-cell RNA-sequencing data from mouse lungs across eight developmental timepoints: E16.5, E18.5, PND1, PND3, PND7, PND10, PND14, and PND28. The dataset was downloaded in H5AD format, which included raw count matrices, cell-level metadata containing cell type annotations (celltype_level3 and celltype_level3_fullname), sample identifiers (DataID), and developmental stage labels (orig.ident). Cell type annotations provided in the H5AD metadata were used directly without additional automated cell type prediction.

Raw count data were imported from the H5AD file and converted to a Seurat object using the anndataR and Seurat R packages (Seurat 5.1.0, RRID:SCR_016341). Cells with fewer than 250 detected features and genes detected in fewer than 30 cells were excluded during object creation using the min.features and min.cells parameters, respectively. Data normalization was performed using Seurat’s NormalizeData function, followed by identification of variable features (FindVariableFeatures), scaling (ScaleData), and dimensionality reduction by principal component analysis (RunPCA, npcs = 50). A shared nearest neighbor graph was constructed using FindNeighbors with the first 30 principal components, and cells were clustered at resolution 0.1 (FindClusters). Two-dimensional embedding was generated using UMAP (RunUMAP, dims = 1:30). Cell types were assigned based on the metadata annotations provided in the LungMAP dataset (celltype_level3_fullname), and major cell populations including AT2 cells, AT1/AT2 transitional cells, and alveolar macrophages were identified and subset for downstream analyses.

For differential expression analysis between developmental stages, pseudobulk count matrices were generated by aggregating raw counts across cells within each biological sample using Seurat’s AggregateExpression function. For AT2 cells, two levels of aggregation were performed: (1) by developmental timepoint (orig.ident) yielding 8 pseudobulk samples, and (2) by individual sample (DataID) yielding 16 pseudobulk samples. Alveolar macrophages were similarly aggregated by individual sample. DESeq2 (version 1.44.0, RRID:SCR_015687) was applied to each pseudobulk count matrix using a single-factor design (~age) comparing embryonic versus postnatal stages. For alveolar macrophages, additional per-timepoint DESeq2 contrasts were constructed, in which each developmental stage was compared against all other stages combined (e.g., E16.5 vs. not-E16.5) to identify timepoint-specific differentially expressed genes. Raw counts were rlog-normalized (regularized log transformation, blind = FALSE) prior to downstream analyses including principal component analysis and correlation calculations. PCA was performed on the rlog-transformed values using the plotPCA function from the DESeq2 package, with percent variance explained reported for the top two principal components. Pairwise Spearman rank correlation coefficients were computed across pseudobulk samples to assess global expression similarity between developmental timepoints.

To identify genes whose expression correlates with *Il34* in AT2 cells, Spearman rank correlation coefficients (ρ) were computed between the rlog-normalized expression of Il34 and each gene across pseudobulk samples. Genes with fewer than 3 overlapping observations were excluded. The resulting correlation coefficients were used to generate a ranked gene list for gene set enrichment analysis (GSEA) using the clusterProfiler R package (version 4.16.0) with the Gene Ontology Biological Process (GO-BP) ontology and the org.Mm.eg.db mouse annotation database. Enrichment was assessed using gseGO with a Benjamini–Hochberg adjusted p-value cutoff of 0.05.

#### Hierarchical gene clustering along developmental time

To identify co-expressed gene modules across developmental timepoints, hierarchical clustering was performed on the rlog-normalized pseudobulk expression matrix. Genes were z-score normalized across samples (row scaling), and genes with zero or undefined variance were removed. Pairwise distances between genes were calculated using Euclidean distance, and agglomerative clustering was performed using Ward’s linkage method (ward.D2). The resulting dendrogram was cut into k = 12 clusters for AT2 cells and k = 2 clusters for alveolar macrophages. Heatmaps were generated using the ComplexHeatmap R package (version 2.20.0, RRID:SCR_017270) with z-score scaled expression values displayed on a blue–white–red color scale (range: −3 to +3). Gene cluster assignments were saved as CSV files and per-cluster gene lists exported as text files for downstream pathway analysis. For AT2 cells, Spearman correlation matrices were computed among genes within selected clusters, filtered at |ρ| > 0.7, and visualized as correlation heatmaps using pheatmap.

To characterize temporal expression dynamics, normalized expression values from pseudobulk-aggregated Seurat objects (LogNormalize) were extracted for select genes of interest. For AT2 cells, normalized expression and percent gene-positive cells were computed across developmental timepoints and individual samples, with means and standard deviations displayed as error bars. Genes of interest included *Il34*, *Csf2*, *Sftpd*, *Gpx1*, *Prdx1*, *Hmga2*, *Cldn4*, *Itga2*, and *Slc4a11*. Developmental timepoints were ordered as E16.5, E18.5, PND1, PND3, PND7, PND10, PND14, and PND28. Individual gene expression plots were generated using ggplot2 and saved as PDFs.

### Pathway enrichment analysis in LungMAP

Gene Ontology (GO) enrichment analysis was performed on individual gene clusters using the clusterProfiler R package with org.Mm.eg.db for gene ID conversion. Enrichment was assessed separately for Biological Process (BP), Molecular Function (MF), and Cellular Component (CC) ontologies, with a Benjamini–Hochberg adjusted p-value cutoff of 0.05. KEGG pathway enrichment was performed using enrichKEGG (organism = mmu) with an adjusted p-value cutoff of 0.05 and q-value cutoff of 0.2, followed by setReadable for conversion to gene symbols. Results were visualized using dot plots and bar plots.

For alveolar macrophages, gene set enrichment analysis (GSEA) was performed using the desktop GSEA application (Broad Institute, RRID:SCR_003199) in pre-ranked mode. Wald statistics from each per-timepoint DESeq2 contrast were exported as .rnk files and used as ranked input gene lists. Enrichment was assessed against the GO Biological Process (GOBP), Hallmark, and KEGG pathway collections from the Molecular Signatures Database (MSigDB, version 2024.1). Normalized enrichment scores (NES) and FDR q-values were extracted from GSEA output reports for both positively and negatively enriched gene sets. GSEA results were visualized as bubble plots showing NES magnitude and direction across developmental timepoints, with gene sets hierarchically clustered by their NES profiles. Significance was annotated by FDR q-value thresholds as indicated in individual figures.

#### Analysis of merged alveolar epithelial and tumor cells scRNA-seq

To perform the merged AT/tumor cell analysis in Fig. 5a-j, AT/tumor cell clusters from the *SftpcCreKras*^G12D^ anti-IL34 vs control and *Il34*^−/−^ vs *Il34*^+/+^ experiments were identified, extracted and merged, followed by re-normalization, clustering and dimensionality reduction using standard Seurat v5 functions. Classification of the new clusters identified non-epithelial cells and doublets which were removed from analysis. Clustering was repeated on the filtered dataset, resulting in 9 clusters and 7,292 cells and comprising the following experimental groups: *SftpcCre* (tumor-free), *Il34*^−/−^ *SftpcCre* (IL34-deficient tumor-free), *SftpcCreKras*^G12D^ anti-IL34 (tumor anti-IL34 treated), *SftpcCreKras*^G12D^ control (tumor control treated), *Il34*^−/−^ *SftpcCreKras*^G12D^ (IL34-deficient tumor), *Il34*^+/+^ *SftpcCreKras*^G12D^ (IL34-wild type tumor). To assess cluster frequencies, the number of cells in each cluster in each sample was divided by the total number of AT/tumor cells in the respective sample. GSEA comparison between *Il34*^−/−^ *SftpcCreKras*^G12D^ and *Il34*^+/+^ *SftpcCreKras*^G12D^ groups was performed using the Java GSEA program in pre-ranked mode on DESeq2 differential expression results obtained using DESeq2 as described previously. DESeq2 results were used to perform overall DEG analysis and consensus DEG analysis.

To calculate RNA velocity and transition probabilities, the output of velocyto was subset to match the cells present in the AT/tumor cell Seurat object in R, then subset to create objects for the *Il34*^−/−^ *SftpcCreKras*^G12D^ and *Il34*^+/+^ *SftpcCreKras*^G12D^ groups as separate objects, exported using the anndataR package and further processed in Python v3.12 using the anndata and scvelo packages. The scvelo vignette was used to select commands to calculate RNA velocity vectors, predicted cluster transition probabilities, pseudo time and partition-based graph abstraction (PAGA) predicted cluster transition directionality.

#### Software and statistical methods

All analyses were performed in R (version 4.5) unless otherwise noted. Key R packages and versions used were Seurat 5.1.0 (RRID:SCR_016341), DESeq2 1.44.0 (RRID:SCR_015687), clusterProfiler 4.16.0, ComplexHeatmap 2.20.0 (RRID:SCR_017270), pheatmap, ggplot2 (RRID:SCR_014601), anndataR, vsn, reshape2, copilot, dplyr, tidyr, readr, tibble, patchwork, ragg, circlize, and cowplot. Gene set enrichment analysis was performed using the GSEA desktop application (RRID:SCR_003199) and the fgsea and clusterProfiler R packages. The Molecular Signatures Database (MSigDB) version 2024.1 was used for all GSEA analyses. Statistical significance for differential expression was defined as adjusted p-value < 0.05. As of 5/14/26 the Python package version used were: anndata 0.11.4, scvelo 0.3.3, numpy 1.26.0, scipy 1.15.2. Plots were generated using ggplot2, ComplexHeatmap, and pheatmap, and saved as PDF files. Microsoft Copilot 365 was used to assist with code troubleshooting.

#### TCGA analyses

To analyze the relationship between individual gene expression and overall survival, TCGA data available through cBioPortal was subjected to survival analysis. The Genomics Data Commons version of each TCGA dataset was used. Unless otherwise stated, the threshold for gene expression was as follows: the mRNA expression z-score>0 was used to identify highly expressing samples and z-score<0 were low expressing samples. The log rank test p-value was reported. For hazard ratios, the low-expressing samples were used as the reference group with hazard ratios reported for high-expressing tumors relative to the low-expressing reference. To perform subgroup analysis by KRAS mutation status, tumor stage and other parameters, GDC data available through cBioPortal or directly through GDC were used to create sample subsets. The cBioPortal release of May 6, 2025, was used. To analyze co-expression, the cBioPortal calculated Spearman rank correlation coefficients were used based on default settings.

### Statistical analysis

Experimental data were graphed and expressed as the mean +/− standard error of the mean (mean +/− SE) using Graph Pad version 10.0 (RRID:SCR_002798). Results were analyzed statistically using one-way ANOVA with Tukey’s post-hoc test for multiple group analysis and Student’s t-test or Mann Whitney test for two-group analysis using GraphPad Prism version 9.1.0. Data with *p*≤0.05 were considered statistically significant and values greater than p>0.05 were considered non-significant. All studies were performed 3 or more times with three or more replicates per group. Statistical analyses of RNA-sequencing and scRNA-sequencing data were performed using R and python packages as described above. R version 4.5 and python version 3.12 were used.

## Data, code and materials availability

All data associated with this study are in the main manuscript or extended data materials. Source data will be available at 10.6084/m9.figshare.30633545 once published and during the review process at https://figshare.com/s/2f8307e851090f663eef. The RNA sequencing data discussed in this publication have been deposited in NCBI’s Gene Expression Omnibus^57^ and are accessible through Gene Expression Omnibus (GEO, RRID:SCR_005012) Series accession numbers as follows: RNA sequencing of sorted LLC macrophages GSE119520, Single cell sequencing of LLC tumors by Rhapsody platform GSE121602, 10X Genomics single cell sequencing of LLC tumors GSE310030 (reviewer token qvuxukgcxlotlov), single cell sequencing of *CC10Cre* vs *CC10Cre;Kras^G12D^;p53^fl/fl^* lung tissue GSE310268 (reviewer token spkxaamczluxpsn), single cell sequencing of *CC10Cre;Kras^G12D^;p53^fl/fl^* lungs +/− anti-IL34 treatment GSE311061 (reviewer token unqzyykkhjizpcz), bulk RNA sequencing of IL-34 with or without anti-IL-34 treated AMs sorted from S*fptcCreKras^G12D^* GSE310269 (reviewer token cjoxcqgipzsdhwx), bulk RNA sequencing of bronchoalveolar lavage cells GSE310270 (reviewer token cpkbwkscthkxxoj), bulk RNA sequencing of *SfptcCre vs SfptcCreKras^G12D^* sorted cells GSE310815 (reviewer token evuxoegehzehhuf), single cell RNA sequencing of *SfptcCreKras^G12D^* vs *SfptcCreKras^G12D^;Il34−/−* lungs GSE311849 (reviewer token ofqreemkjpyxryp), bulk sequencing of *SfptcCreKras^G12D^* vs *SfptcCreKras^G12D^*;*Il34−/−* AMs and tumor cells GSE311062 (reviewer token ctqxucgivvqdxwr), and single cell sequencing of *SftpcCreKras^G12D^* lungs +/− anti-IL34 treatment GSE311059 (reviewer token gpyfwucynpclpaf). Resources are available upon reasonable request to J.A.V.

## Funding

This study was supported by National Institutes of Health grants R01CA167426 and a Targeted Research grant from the Curebound Foundation to J.A.V. as well as a Cancer Research Institute/Irvington postdoctoral fellowship to J.Z. J.Z. is supported by the Huntsman Cancer Institute/Huntsman Cancer Foundation and the National Institutes of Health grant K22CA292568. This work includes data generated at the UCSD Moores Cancer Center using a 10X Genomics Chromium Connect, which was purchased with support of NIH S10OD032398 and at the UCSD IGM Genomics Center using a NovaSeq6000, which was purchased with the support of NIH S10OD026929. This study was also supported by the UCSD Centers for Computational Biology & Bioinformatics Shared Resource and the Tissue Technology Shared Resource of the UCSD Moores Comprehensive Cancer Center supported by National Cancer Institute Cancer Center Support Grant P30CA23100.

## Author contributions

This study was conceptualized and supervised by J.A.V., M.O. and J.Z. Animal models of lung cancer were developed and performed by H.C., G.M., P.C., M.P., Z.X., M.J.H. and J.R. Human and murine lung tissue specimens were collected and provided by P.C. and M.O. J.H. provided clinical pathology review of human lung cancer specimens. H.C. and G.M. performed flow cytometry and data analysis. H.C. and Z.X. performed cell proliferation, Western blotting and ELISA assays. A.R., M.P., A.G., M.J.H. and B.L. performed histological analyses. E.W. and M.P. performed single cell sequencing. J.Z., P.O., E.W., and R.S performed bioinformatic analyses. J.Z. and P.O. performed software development and database mining. M.C. provided IL34−/− mice. H.F., M.De P., M.P. and M.M. provided single cell sequencing datasets. All authors analyzed data and reviewed and approved the manuscript.

## Competing interests

J.A.V. is a founder of Javelin Biomedicine and a shareholder in AlphaBeta Therapeutics and EMT Biosciences. The other authors declare no competing interests.

## Extended Data

**Extended Data Figure 1:**
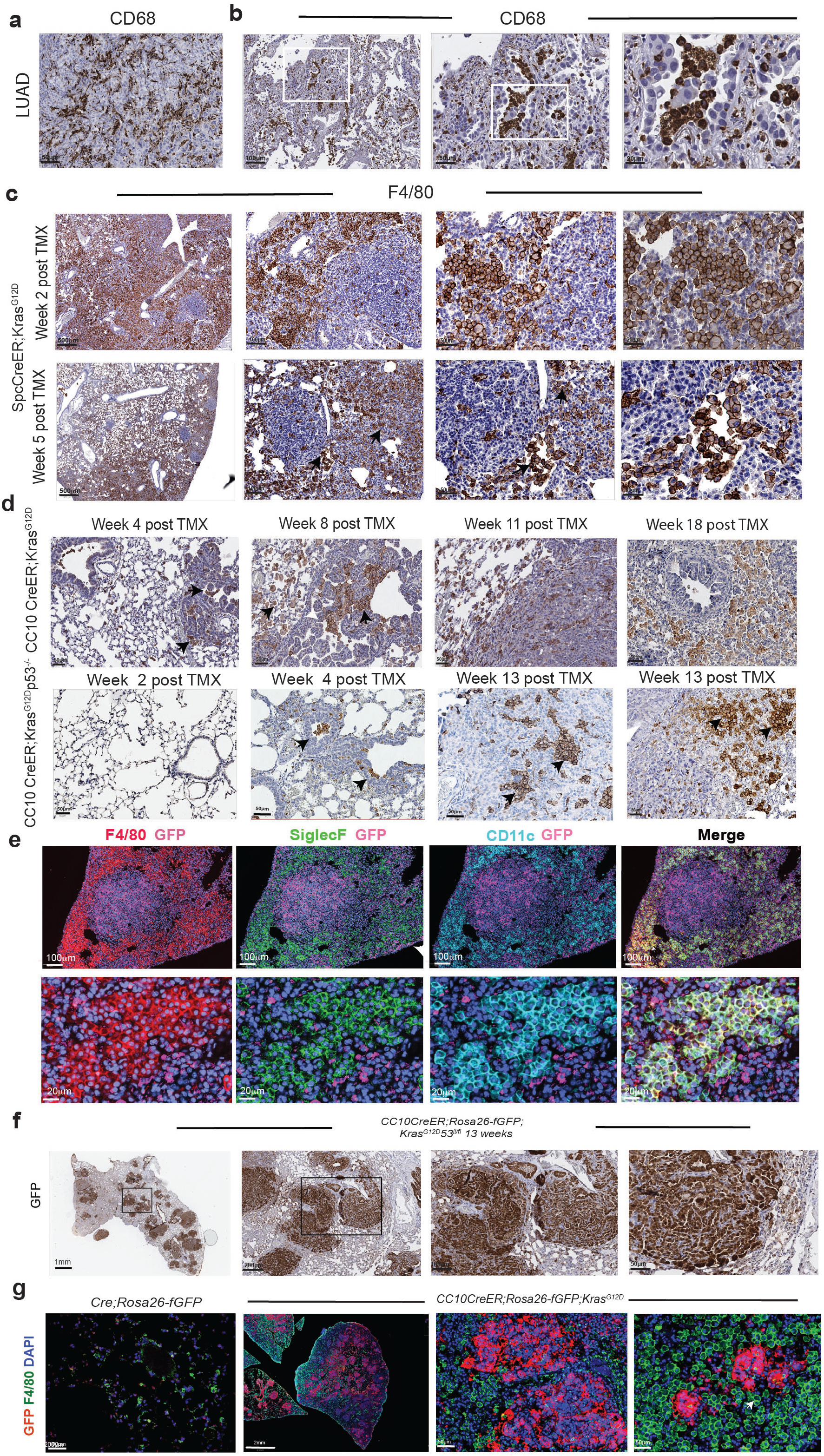
Histological analysis of murine KrasG12D mutant lung tumors reveals alveolar macrophage expansion during tumor progression. (a-b) Fusiform (a) and alveolar (b) CD68+ macrophages in stage lung 1 adenocarcinomas (LUAD) at successive magnifications. (c) F4/80+ macrophages in lungs from *SftpcCreER;Kras^G12D^* animals at two and five weeks post-tamoxifen administration at successive magnifications. (d) F4/80+ macrophages in lungs of *CC10CreER;Kras^G12D^ and CC10CreER;Kras^G12D^p53^fl/fl^* animals at various times post-tamoxifen administration. (e) F4/80+ (red), SiglecF+ (green), CD11c+ (cyan) and GFP+ (pink) immunostaining in lungs from *SftpcCreER;Rosa26-fGFP;LSL-Kras^G12D^* mice five weeks post-tamoxifen administration. (f) GFP+ tumor nodules at successive magnifications in lung tissue from an *CC10Cre;Rosa26-fGFP;Kras^G12D^p53^fl/fl^* animal 13 weeks post-tamoxifen treatment. (g) Anti-GFP (red), anti-F4/80 (green) and Dapi (blue) immunofluorescence in lungs from *CC10Cre;Rosa26-fGFP;Kras^G12D^* mice 11 weeks post tamoxifen.

**Extended Data Figure 2:**
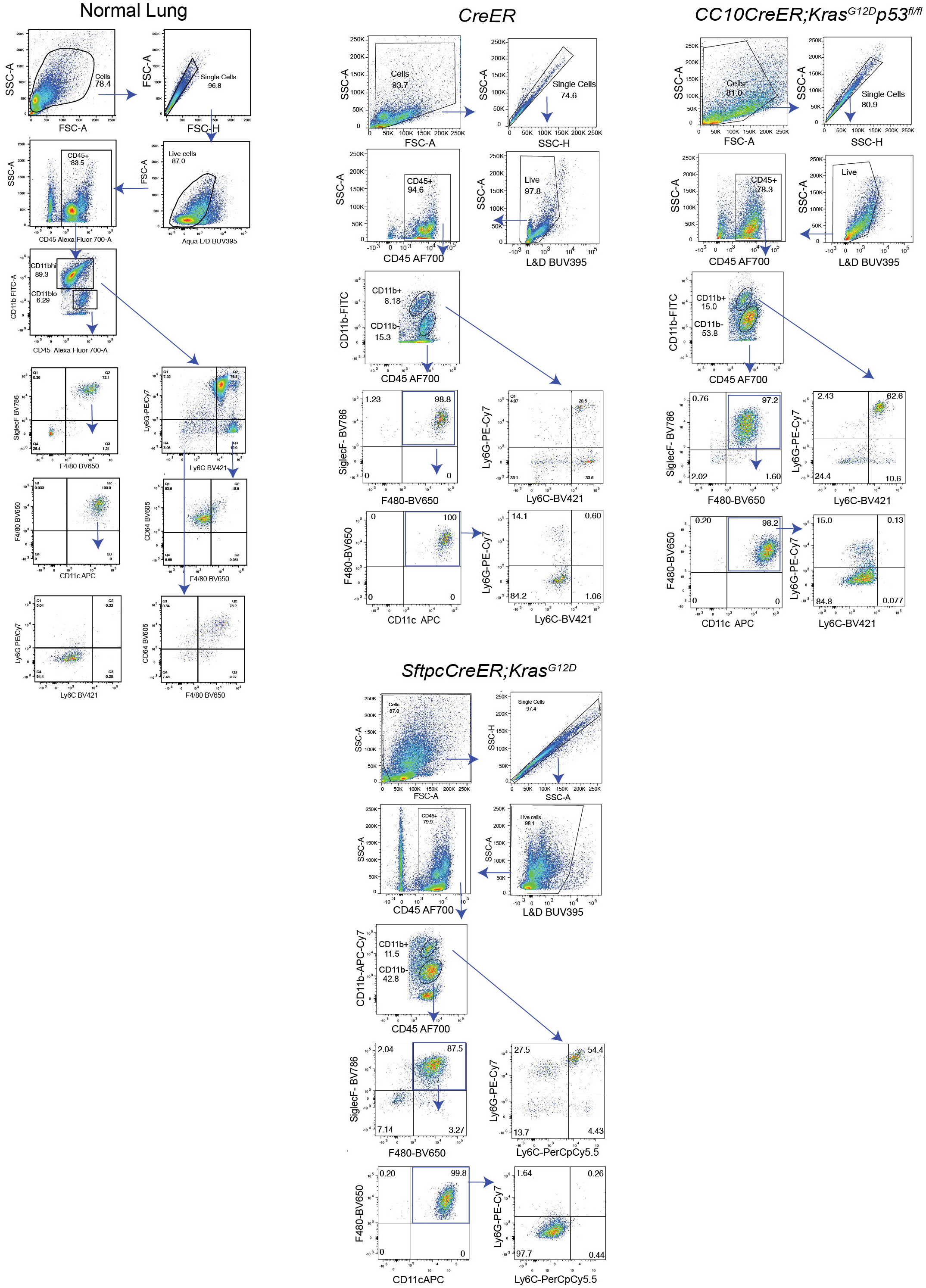
Flow cytometry gating. Flow cytometry gating scheme to detect myeloid cells (CD11b-SiglecF+F4/80+ CD11c+alveolar macrophages, CD11b+Ly6C+Ly6G-monocytes, CD11b+Ly6C-Ly6G-F4/80hi interstitial macrophages and CD11b+ Ly6G+Ly6C+neutrophils) in lungs from *C57BL6* (normal lung), *CC10Cre, CC10CreER;LSL-Kras^G12D^p53^fl/fl^* and *SftpcCreER;LSL-Kras^G12D^* animals.

**Extended Data Figure 3:**
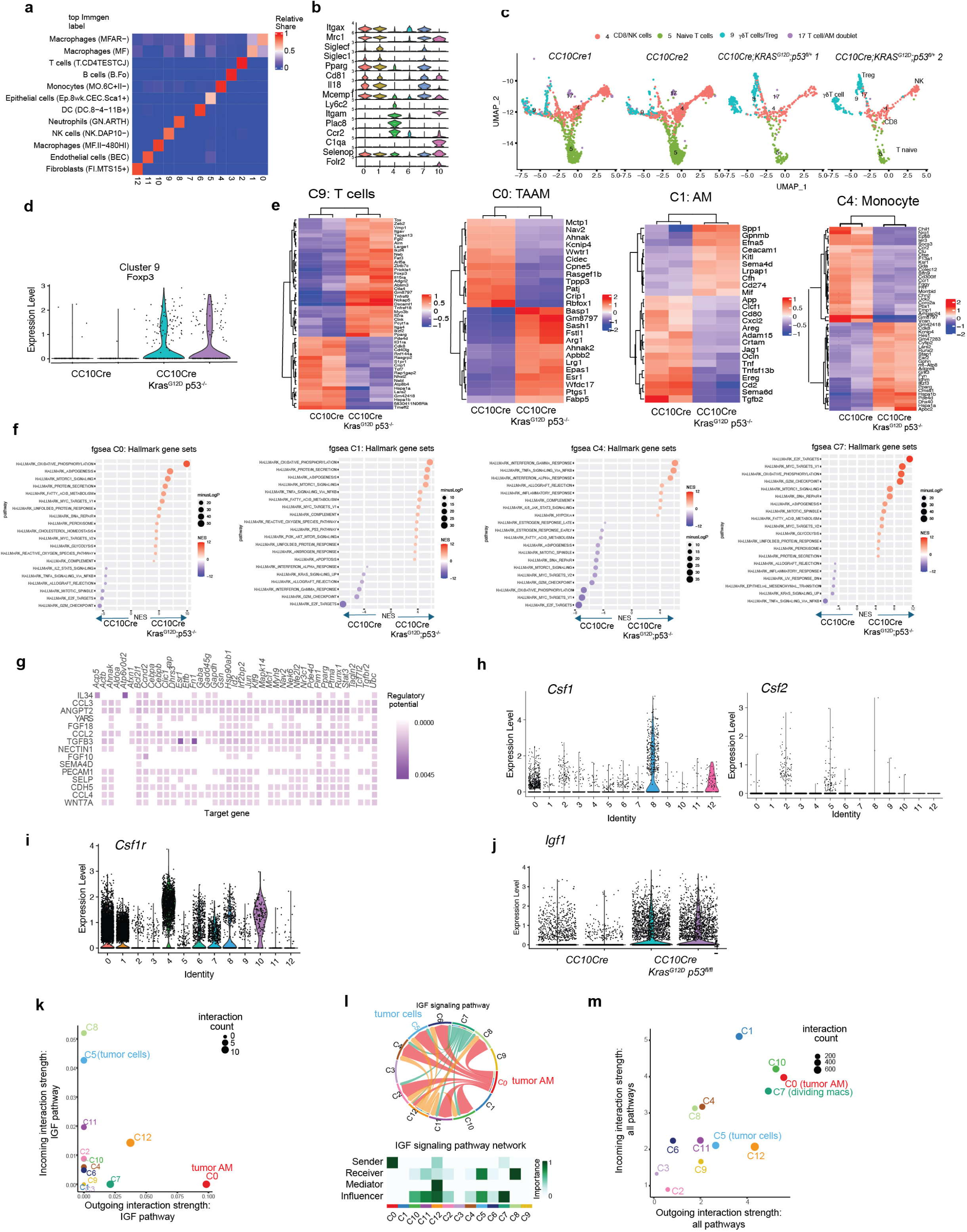
*CC10CreER;Kras^G12D^p53^fl/fl^* model immune profile by scRNA sequencing. (a) Heatmap of top Immgen category scores in single cell sequencing of lung samples from *CC10Cre* and *CC10CreER;Kras^G12D^;p53^fl/fl^* animals as determined by SingleR. (b) Violin plots of myeloid cell markers expressed in single cell clusters from *CC10Cre* and *CC10CreER;Kras^G12D^;p53^fl/fl^* animals. (c) T cell clusters in *CC10CreER* and *CC10CreER;Kras^G12D^p53^fl/fl^* samples. (d) Violin plot depicting Fox P3 expression in cluster 9 T cells. (e) Heatmap of top differentially expressed genes in T cells from cluster 9 and of top differentially expressed genes in clusters C0, C1, and C4. (f) Expression of GSEA Hallmark signatures in Clusters C0, C1, C4 and C7. (g) Target gene signatures from Cytosig analysis of interactions between cluster C0 and C5. (h) Violin plots of *Csf1* and *Csf2* expression in clusters from Figure 2a. (i) Violin plot of *Csf1r* expression in clusters from 2a. (j) Violin plot of *Igf1* expression in Clusters C0 and C1 in *CC10CreER* and *CC10CreER;Kras^G12D^p53^fl/fl^* samples. (k) Incoming and outgoing interaction strength of the IGF pathway in each cluster in samples as determined by CellChat v2. (l) Chord plot depicting differential interaction strengths between clusters as determined by CellChat v2. (m) Total Incoming and outgoing strength of all annotated pathways in lung samples from *CC10Cre* and *CC10CreER;Kras^G12D^;p53^fl/fl^* animals.

**Extended Data Figure 4:**
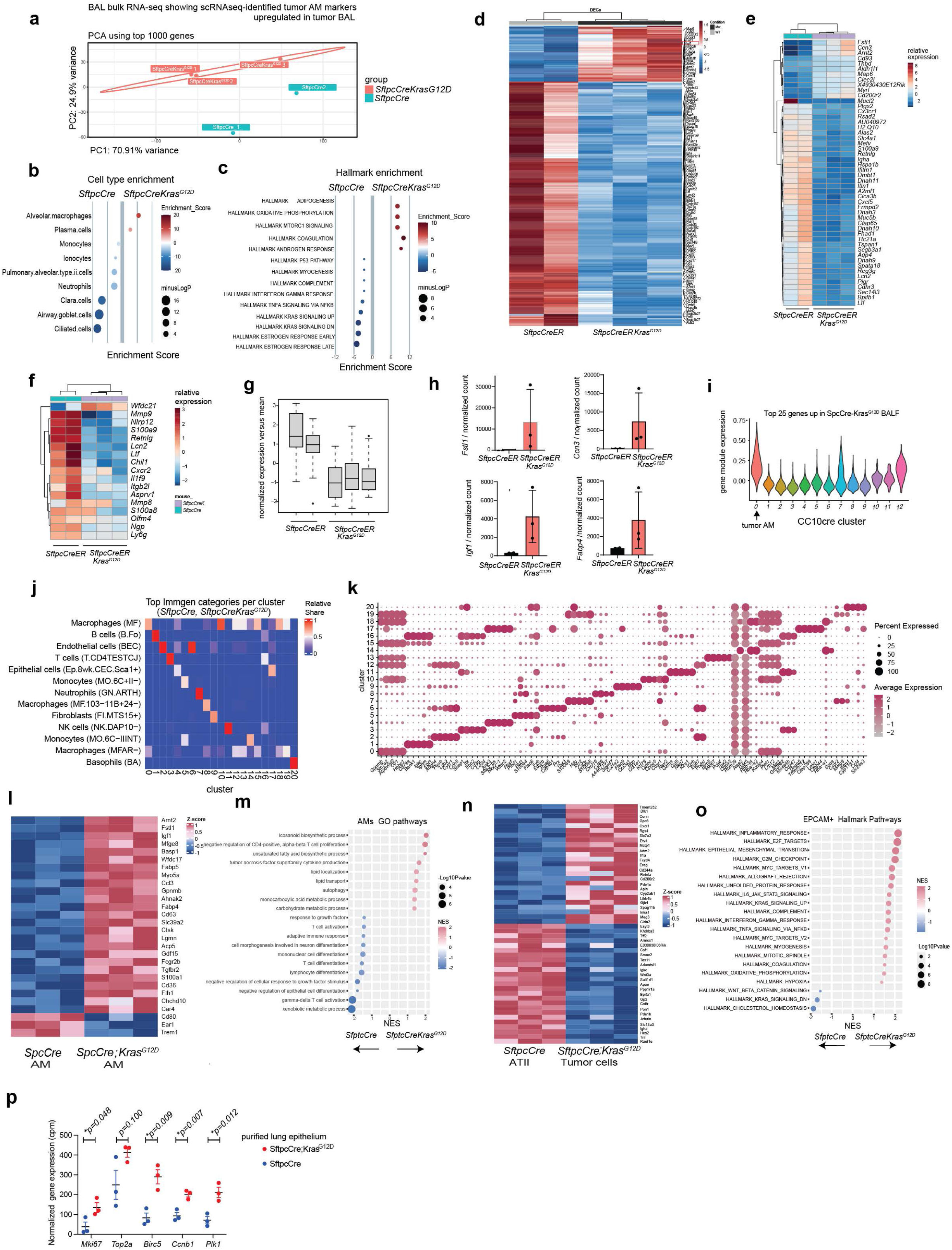
Characterization of alveolar macrophages in *SftpcCre* and *SftpcCreER;KrasG12Danimals*. (a) Principal component map of RNAs sequences of cells from bronchoalveolar lavage of lungs from 12 week old *SftpcCre* (n=2) and *SftpcCreER;Kras^G12D^* (n=3) mice. (b-c) Plots of differential (b) cell type and (c) Hallmark pathway enrichment in BAL cells from *SftpcCre* and *SftpcCreER;Kras^G12D^*. (d-e) Heatmaps of (d) top 100 differentially expressed genes and (e) the top 50 differentially expressed gene in BAL cells. (f) Differentially expressed neutrophil genes in BAL cells from *SftpcCre* and *SftpcCreER;Kras^G12D^*. (g) Box and whisker plot of normalized counts of 20 neutrophil genes from (f). (h) Normalized counts of tumor alveolar macrophage genes in BAL cells from *SftpcCre* and *SftpcCreER;Kras^G12D^* animals. (i) Violin plot of expression of top 25 upregulated genes from *SftpcCreER;Kras^G12D^* BAL cells in *CC10CreKras^G12D^p53^fl/fl^* clusters. (j) Heatmap of top Immgen category scores and (k) dot plot of top cluster markers in single cell sequencing of lung samples from *SftpcCre* and *SftpcCreER;Kras^G12D^* animals. (l) Heatmap of differential expression of AM genes in sorted alveolar macrophages from lungs of *SftpcCre* and *SftpcCreER;Kras^G12D^* animals. (m) GSEA GO pathway expression in purified alveolar macrophages from *SftpcCre* and *SftpcCreER;Kras^G12D^* lungs. (n) Heatmap of differential expression in purified epithelial cell/tumor cells from *SftpcCre* and *SftpcCreER;Kras^G12D^* lungs. (o) GSEA Hallmark pathway expression in bulk sequencing from purified epithelial cell/tumor cells from *SftpcCre* and *SftpcCreER;Kras^G12D^* lungs. (p) Normalized expression of proliferation genes in purified lung epithelium/tumor cells from *SftpcCre* (n=3) and *SftpcCreER;Kras^G12D^* lungs (n=3).

**Extended Data Figure 5:**
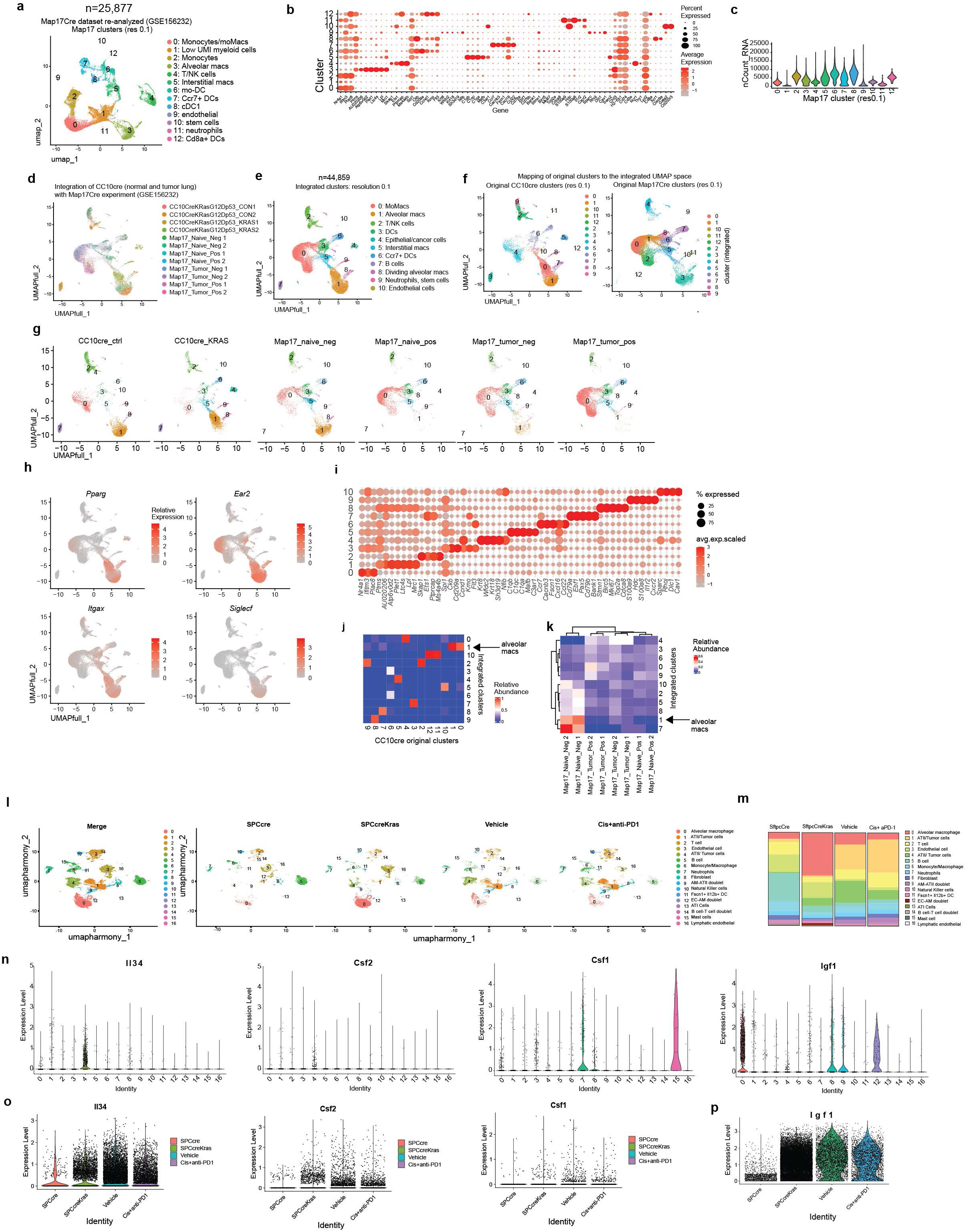
Analysis of anti-IL34 treated *SftpcCre;Kras^G12D^* tumors. (a) Mean fluorescence intensity of CSF1R and CSF2R expression on alveolar macrophages and CD11b+Ly6C+ monocyte/macrophages from *SfptcCre and SfptcCre;Kras^G12D^* mice. (b) Western blot depicting time course of CSF1R, ERK and AKT phosphorylation upon stimulation of mouse bone marrow derived macrophages with mCSF or IL-34 (50ng/ml). (c) Siglec F+ F4/80+ CD11c+ alveolar macrophage content determined by flow cytometry in lungs from *SfptcCre; SfptcCre;Kras^G12D^ and SfptcCre;Kras^G12D^;Il34−/−* mice. (d) Percent CD4+ and CD8+ T cells in *SfptcCre;Kras^G12D^* mice treated with anti-IL-34 (n=4) or isotype control (n=4). (e) Images and graph of Siglec F+ alveolar macrophages/mm^2^ in lungs from *SfptcCre;Kras^G12D^*mice treated with anti-IL34 (n=3) or isotype control (n=5) or from *SfptcCre;Kras^G12D^;Il34−/−* mice (n=4). (f-g) Merged (f) and groupwise (g) UMAP plots of cell populations of integrated scRNA-sequencing samples from lungs of control and anti-IL34 treated *SftpcCreER;Kras^G12D^* animals (n=3-4 replicates per genotype). (h) Dot plot of selected marker genes of the dataset. (i) Color-coded fold-changes in the proportion of cells in clusters from anti-IL34 treated *SftpcCreER;Kras^G12D^* lungs relative to control treated *SftpcCreER;Kras^G12D^* lungs. (j) Proportions of cells in cluster 0 and 1 in control and anti-IL34 treated *SftpcCreER;Kras^G12D^*samples. (k-l) Dot and violin plots of (k) tumor alveolar macrophages (TAAM) and (l) normal AM markers expressed in clusters 0 and 1 from (f). (m-o) Heat maps of differential gene expression in (m) C0, (n) C1 and (o) C6 from (f). (p-s) GSEA Hallmark pathway expression in Cluster 0 and 1 from (f).

**Extended Data Figure 6:**
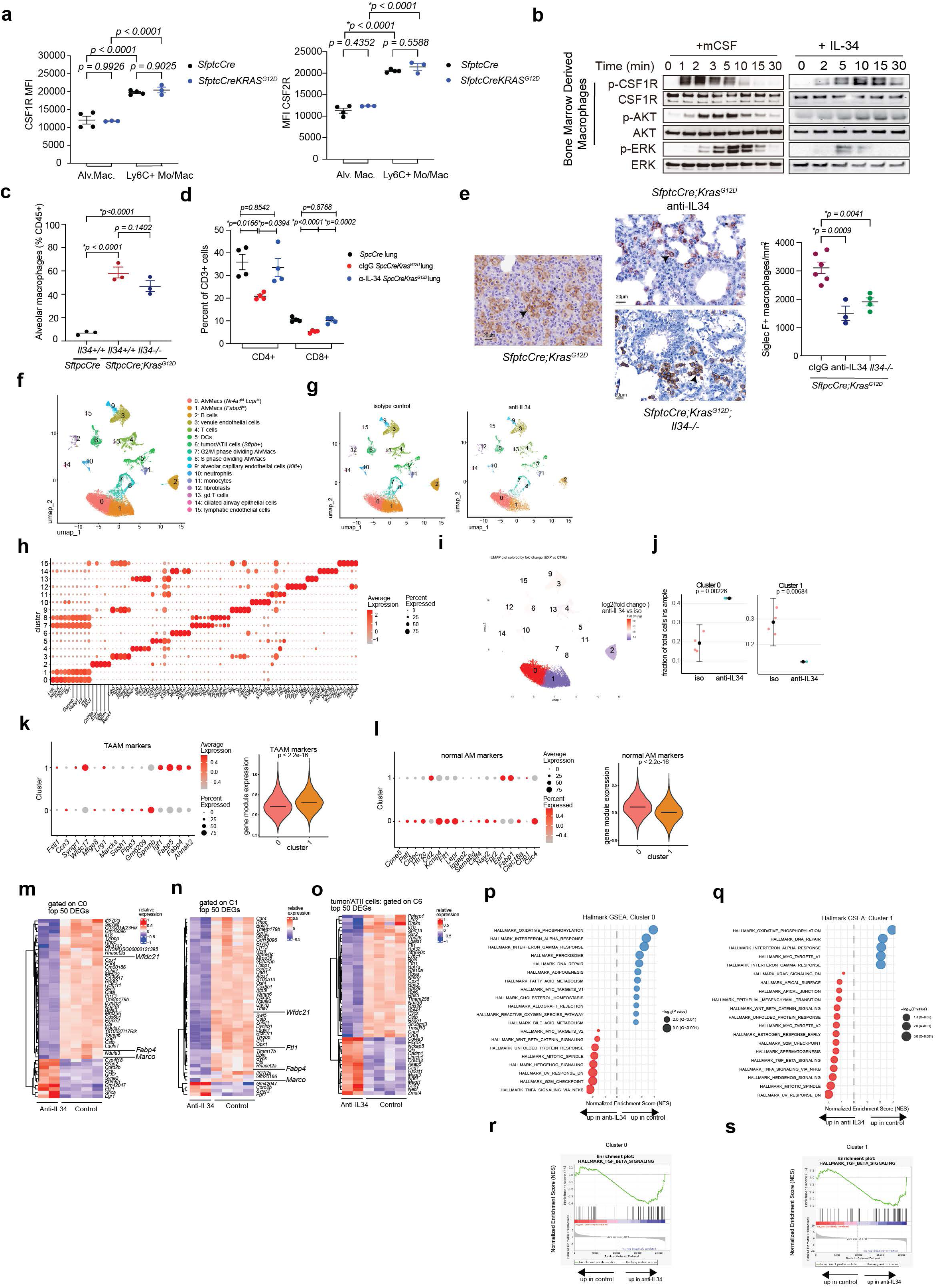
Analysis of anti-IL34 treated *SftpcCre;Kras^G12D^* tumors. (a) Mean fluorescence intensity of CSF1R expression on alveolar macrophages and CD11b+Ly6C+ monocyte/macrophages from *SfptcCre and SfptcCre;Kras^G12D^* mice. (b) Western blot depicting time course of CSF1R, ERK and AKT phosphorylation upon stimulation of mouse bone marrow derived macrophages with mCSF or IL-34 (50ng/ml). (c) Siglec F+ F4/80+ CD11c+ alveolar macrophage content determined by flow cytometry in lungs from *SfptcCre; SfptcCre;Kras^G12D^ and SfptcCre;Kras^G12D^;Il34−/−* mice. (d) Percent CD4+ and CD8+ T cells in *SfptcCre;Kras^G12D^* mice treated with anti-IL-34 (n=4) or isotype control (n=4). (e) Images and graph of Siglec F+ alveolar macrophages/mm^2^ in lungs from *SfptcCre;Kras^G12D^* mice treated with anti-IL34 (n=3) or isotype control (n=5) or from *SfptcCre;Kras^G12D^;Il34−/−* mice (n=4). (f-g) Merged (f) and groupwise (g) UMAP plots of cell populations of integrated scRNA-sequencing samples from lungs of control and anti-IL34 treated *SftpcCreER;Kras^G12D^* animals (n=3-4 replicates per genotype). (h) Dot plot of selected marker genes of the dataset. (i) Color-coded fold-changes in the proportion of cells in clusters from anti-IL34 treated *SftpcCreER;Kras^G12D^* lungs relative to control treated *SftpcCreER;Kras^G12D^* lungs. (j) Proportions of cells in cluster 0 and 1 in control and anti-IL34 treated *SftpcCreER;Kras^G12D^* samples. (k-l) Dot and violin plots of (k) tumor alveolar macrophages (TAAM) and (l) normal AM markers expressed in clusters 0 and 1 from (f). (m-o) Heat maps of differential gene expression in (m) C0, (n) C1 and (o) C6 from (f). (p-s) GSEA Hallmark pathway expression in Cluster 0 and 1 from (f).

**Extended Data Figure 7:**
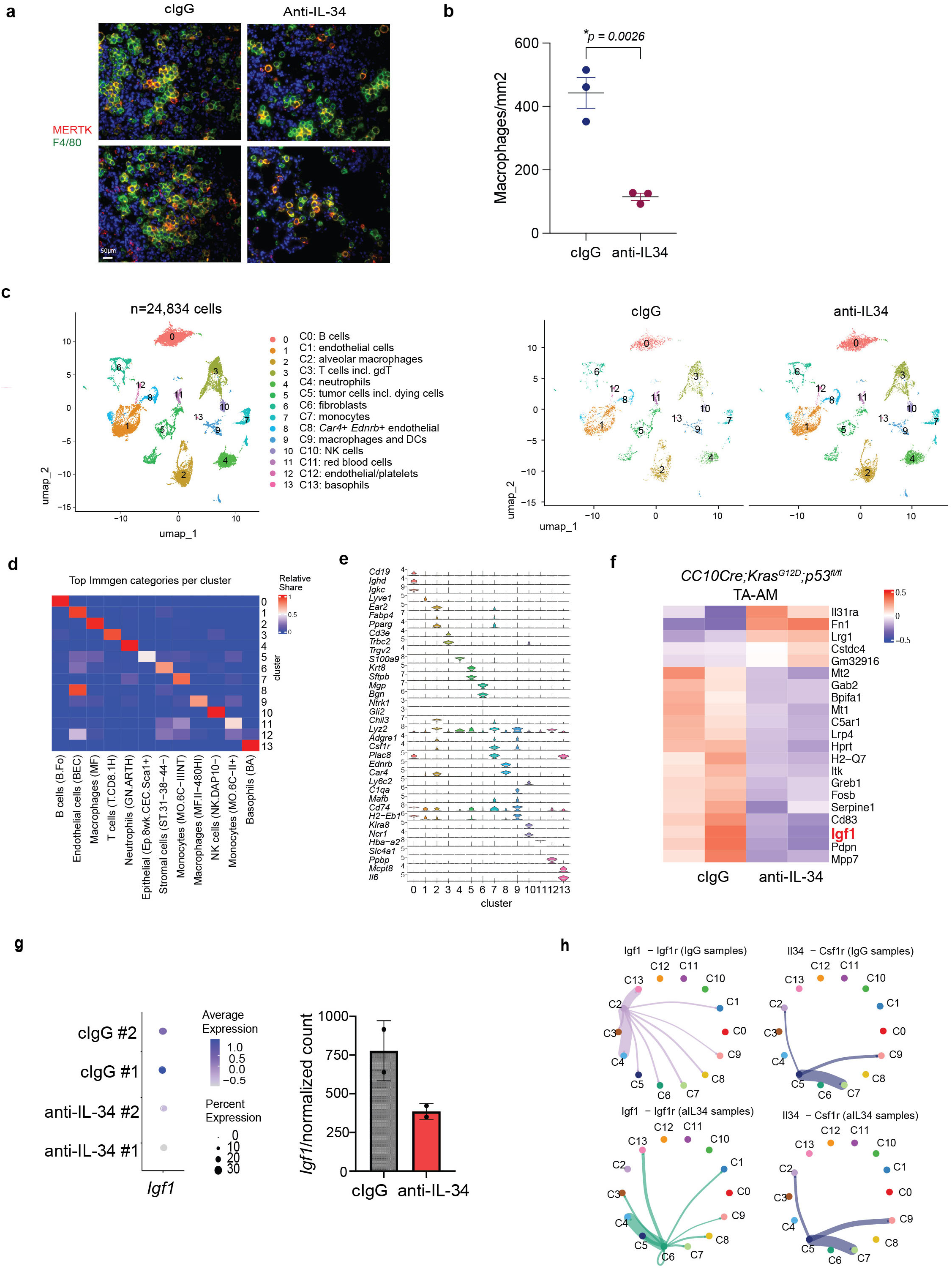
Analysis of anti-IL34 treated *CC10Kras^G12D^;p53^fl/fl^* tumors. (a) F4/80 (green) and MerTK (red) immunostaining of anti-IL34 or control treated *CC10CreER;Kras^G12D^p53^fl/+^* lungs. Yellow indicates overlap of markers. (b) Mean +/−SEM MerTK+ F4/80+ cells from a (n=3). (c) Merged and groupwise UMAPs of anti-IL34 or control treated *CC10CreER;Kras^G12D^p53^fl/fl^* scRNA sequencing lung samples. (d) Heatmap of top Immgen category scores and (e) violin plot of myeloid cell genes in anti-IL34 or control treated *CC10CreER;Kras^G12D^p53^fl/f^* scRNA sequencing lung samples. (f) Heatmap of differentially expressed genes in TA-AMs from anti-IL-34- vs cIgG (control)-treated *CC10CreKras^G12D^p53^fl/-^* animals. (g) Dot plot and normalized counts of *Igf1* expression in TA-AMs from cIgG and anti-IL-34 treated *CC10CreKras^G12D^p53^fl/fl^* animals. (h) CellChat network plots showing *Igf1-Igf1r* and *Il34-Csf1r* signaling in clusters from cIgG and anti-IL-34-treated samples.

**Extended Data Figure 8:**
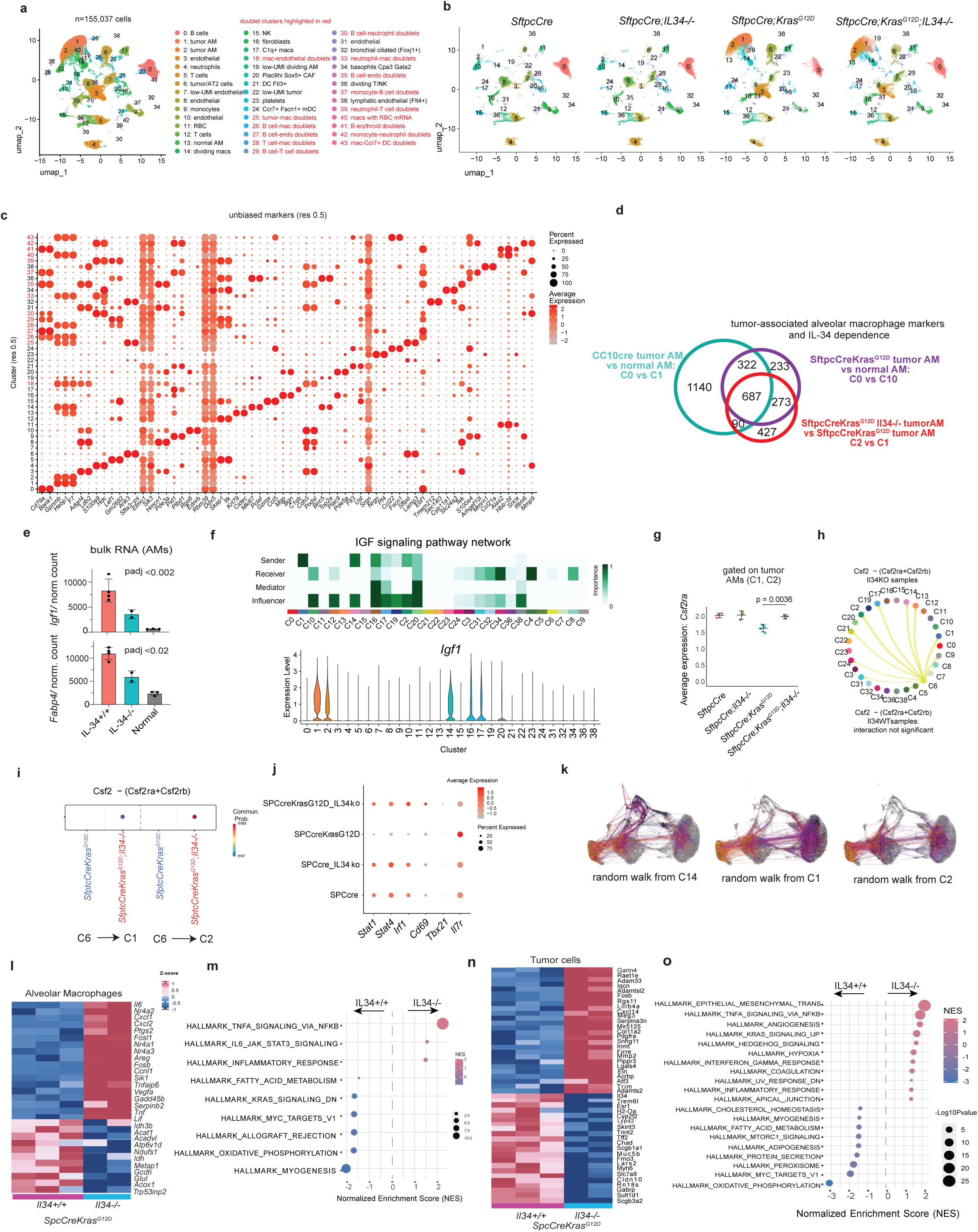
scRNA sequencing analysis of *SftpcCreKras^G12D^; IL34−/−* mouse lungs. (a-b) Merged (a) and groupwise (b) umaps of single cell sequencing of lungs from *SftpcCre, SftpcCre;Il34−/−, SftpcCre;Kras^G12D^,* and *SftpcCre;KrasG12D;Il34*−/− animals (n=3). Doublets are included in a. (c) Dot plots showing expression of population-defining markers from a. (d) Venn diagram depicting shared marker genes in alveolar macrophages from *CC10Cre;Kras^G12D^p53^fl/fl^*, *SftpcCre;Kras^G12D^* and their overlap with alveolar macrophage genes dependent on IL-34 (downregulated in *SftpcCre;KrasG12D;II34*−/− vs *Il34*+/+ alveolar macrophages). (e) *Igf1* and *Fabp4* expression in bulk sequences of *SftpcCre, SftpcCre;Il34−/−, SftpcCre;Kras^G12D^,* and *SftpcCre;KrasG12D;Il34*−/− animals (n=3) samples. (f) IGF-1 signaling network and *Igf1* violin plots in *CC10Cre;Kras^G12D^p53^fl/fl^*, *SftpcCre;Kras^G12D^* and *SftpcCre;KrasG12D;II34*−/− samples. (g) Graph of *Csf2ra* expression per group. (h) CellChat network plot of Csf2-Csf2ra+b signaling in *SftpcCre; Kras^G12D^;Il34−/−*samples (no Csf2 signaling was observed in *SftpcCre;Kras^G12D^ samples*); Cluster 6 (tumor cells). (i) Cell chat signaling probability quantification of the *Csf2-Csf2ra/rb* pathway from Cluster 6 to AM clusters 1 and 2 in *SftpcCre, SftpcCre;Il34−/−, SftpcCre;Kras^G12D^,* and *SftpcCre; Kras^G12D^;Il34−/−*samples. (j) Dot plots of T cell activation genes in T cells from *SftpcCre, SftpcCre;Il34−/−, SftpcCre;Kras^G12D^,* and *SftpcCreIl34−/−Kras^G12D^* samples. Dot plots of markers in T cells from (a). (k) Random walk analysis of RNA velocity quantified by Cellrank 2; black dots indicate points of origin and yellow dots indicate target states. 3 random walk simulations for 3 clusters of origin (C1, C2 and C14) are shown. (l-m) Heat map of differential gene expression (l) and Hallmark Pathway enrichment (m) from bulk sequencing of *SftpcCre;Kras^G12D^* and *StpcCre;Kras^G12D^;Il34−/−* sorted alveolar macrophages. (n-o) Heat map of differential gene expression (n) and Hallmark Pathway enrichment (o) from bulk sequencing of *SftpcCre;Kras^G12D^* and *SftpcCre;KrasG12D;Il34*−/− sorted epithelial/tumor cells.

**Extended Data Figure 9:**
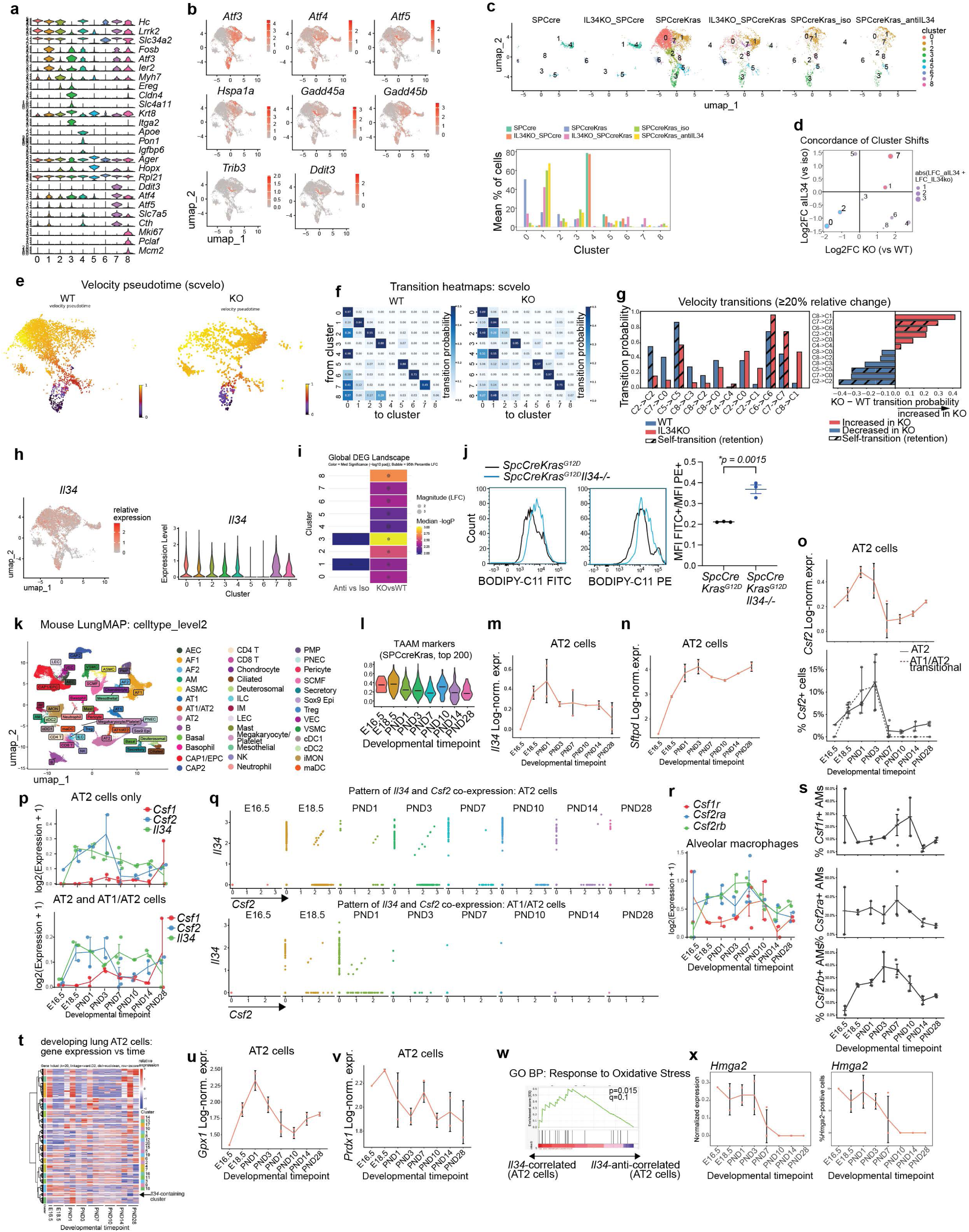
Effect of IL34/IGF1 on ATII cells and alveolar macrophages in KrasG12D tumors and embryonic lung. (a-i) scRNA-seq analysis of merged AT2 and AT1-like cells from *Il34*^−/−^ and anti-IL34 *SftpcCreKras*^G12D^ experiments: (a) violin plot of cluster marker genes; (b) feature plot of cell stress and pre-ferroptosis markers; (c) mean cluster fractions per experimental group. (d) Concordance analysis plot of cluster fraction fold change in *Il34*^−/−^ vs *Il34*^+/+^ tumors (horizontal axis) and anti-IL34 vs control-treated tumors (vertical axis). (e) RNA velocity pseudotime plots for *Il34*^−/−^ vs *Il34*^+/+^ groups predicted by scvelo. (f) Heatmaps of cluster transition probabilities predicted by scvelo. (g) Differences in cluster transition between *Il34*^−/−^ vs *Il34*^+/+^ groups. (h) Feature and violin plots of *Il34* expression in tumor cell clusters. (i) Summary of global differential gene expression changes in clusters assessed by median log2(fold change) magnitude and median −log(p value) for the *Il34*^−/−^ vs *Il34*^+/+^ and anti-IL34 vs control comparisons. (j) Oxidized lipid measurement of *Il34*^−/−^ and *Il34*^+/+^ tumor cells: flow cytometry plots and ratios for oxidized (FITC) and total (PE) lipids shown. (k-x) scRNA-seq analysis of the developing mouse lung based on the LungMAP dataset^42^ (LMEX0000004397): (k) dimensionality reduction and cell type labeling; (l) Violin plot of relative expression of top 200 TAAM markers from the *SPCcreKras*^G12D^ model; (m-o) expression levels of *Il34*, *Sftpd* and *Csf2* in AT2 cells calculated as log of pseudobulk value per biological sample, grouped by developmental timepoint, or as percentage of Csf2+ cells (o bottom panel); (p) expression levels of *Csf1*, *Csf2* and *Il34* in AT1/AT2 transitional and AT2 cells combined (left) and AT2 cells only (right), measured as mean log2(expression+1) per cell; (q) pattern of *Il34* vs *Csf2* expression in AT2 cells (top) and AT1/AT2 transitional cells (bottom). Each dot represents one cell, normalized expression values per cell are shown; (r) expression levels of *Csf1r*, *Csf2ra* and *Csf2rb* in alveolar macrophages, measured as mean log2(expression+1) per cell; (s) expression of *Csf1r*, *Csf2ra* and *Csf2rb* in alveolar macrophages measured as % of transcript-positive cells; (t) heatmap of pseudobulk gene expression values of transcripts expressed in AT2 cells ordered by developmental timepoint and hierarchically clustered by gene. *Il34*-containing cluster is highlighted by an arrow; (u-v) expression levels of *Gpx1* and *Prdx1* in AT2 cells calculated as log of pseudobulk value per biological sample, grouped by developmental timepoint; (w) gene set enrichment plot of the Gene Ontology Biological Process: Response to Oxidative Stress gene set in genes temporally correlated vs anti-correlated with *Il34* expression in AT2 cells; (x) expression levels of *Hmga2* in AT1/AT2 transitional cells measured by mean pseudobulk log expression (left) and as % of transcript-positive cells (right). Statistical comparisons were performed by DESeq2 on pseudobulk counts (f). Error bars show mean ± s.e.m.

**Extended Data Figure 10:**
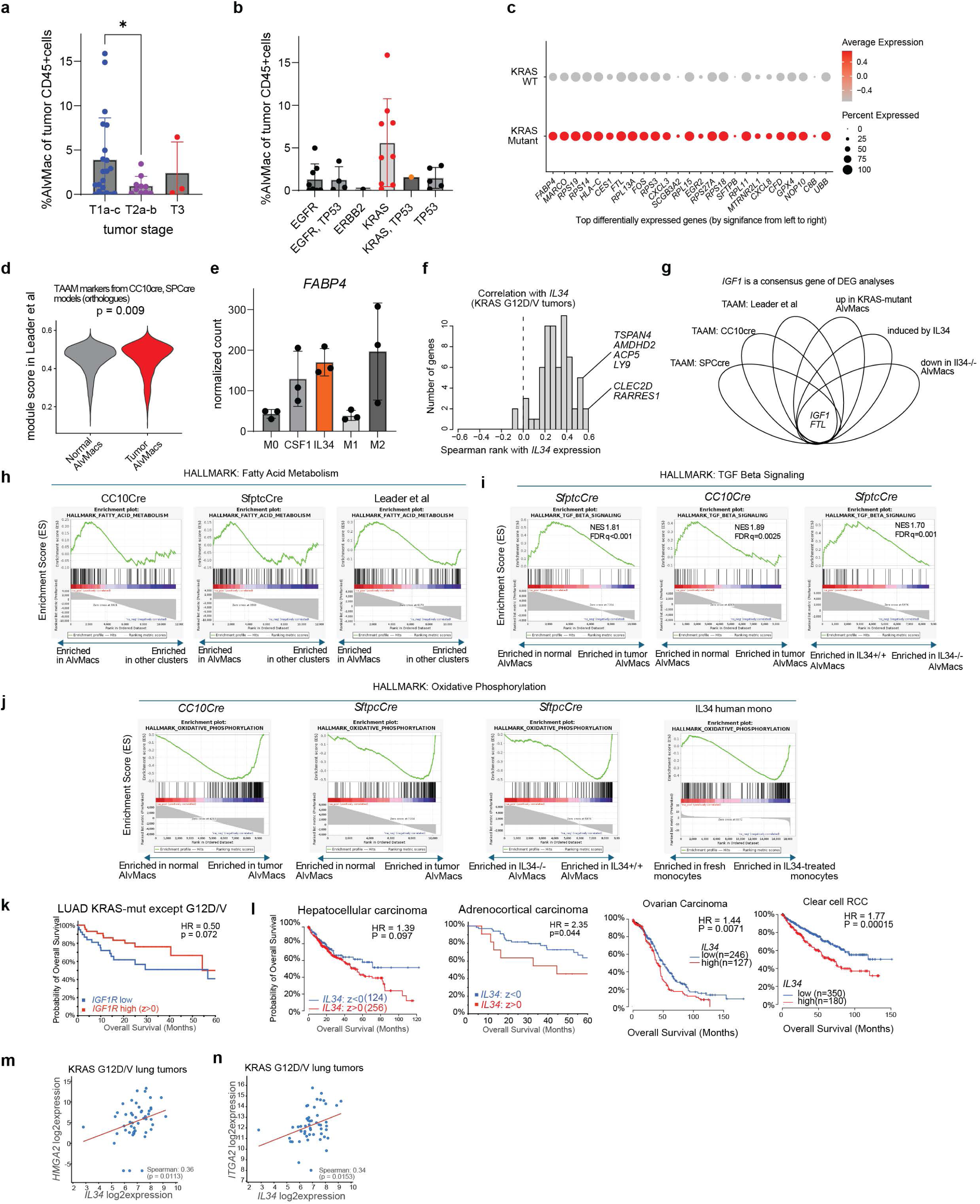
Human resident macrophage RNA sequence analysis. (a-b) Quantification of the percentage of alveolar macrophages of tumor-infiltrating CD45+ cells grouped by tumor stage (a) and tumor somatic mutation (b) in the Leader et al. scRNAseq dataset (GSE154826). (c) Differential expression analysis of alveolar macrophage markers in KRAS-WT vs KRAS-mutant tumors in the GSE154826 dataset. Top 25 genes ranked by p-value are shown. (d) Quantification of tumor-associated alveolar macrophage markers in normal tissue and tumor tissue alveolar macrophages in the GSE154826 dataset. The TAAM markers were derived from the *CC10Cre* and *SftpcCre* GEMMs by consensus analysis of tumor vs normal alveolar macrophage differential expression and obtaining human orthologs of those TAAM genes. (e) Expression of *FABP4* in the in vitro monocyte/macrophage treatment dataset (GSE151194) grouped by treatment group. Normalized counts are shown. (f) Distribution of the Spearman rank correlations of TAAM markers with *IL34* expression in TCGA LUAD KRAS G12D/V tumors. (g) Diagram of the consensus analysis of 6 differential expression comparisons (tumor vs normal AMs *SftpcCre*, tumor vs normal AMs *CC10Cre*, tumor vs normal AMs GSE154826, KRAS-mutant vs KRAS-wild type GSE154826, induced by IL34 in GSE151194 and reduced in Il34−/− vs Il34+/+ AlvMacs in the *SftpcCre* model) depicting *IGF1* and *FTL* as the only consensus genes. (h-j) Consensus analysis of gene set enrichment analyses in the 6 differential comparisons depicted in (f); selected overlapping gene sets are shown. (k) Survival analysis of *IGF1R* expression in KRAS-mutant TCGA LUAD tumors excluding G12D and G12V mutations. (l) Overall survival analysis of IL34 expression in solid tumor TCGA cohorts. (m-n) Correlation between the mRNA levels of *IL34* and HPCS markers *HMGA2* (m) and *ITGA2* (n) in the TCGA lung adenocarcinoma tumors with KRAS G12D or V mutations. Data source is cBioPortal, GDC release of TCGA data. TAAM, tumor-associated alveolar macrophage; TCGA, The Cancer Genome Atlas, GEO, LUAD, lung adenocarcinoma; AM, alveolar macrophage.

